# The collapse indicators of the dominant species outperform community-level critical slowing down indicators

**DOI:** 10.1101/2020.04.27.063784

**Authors:** Melinda Pálinkás, Levente Hufnagel

## Abstract

We studied similarities of collapses and concluded that dominant species have a universal effect on the pattern of collapses. We used open paleoecological and modern data to detect ‘early warning signals’ of collapses. We tested and ranked collapse indicators at the community level (abundance, species richness, constancy, dominance, Shannon’s H, standard deviation, variance, lag-1 autocorrelation of community abundance, lag-1 autocorrelation of Shannon’s H) and at the level of the dominant species (total changes of pre-collapse and collapse dominant species, lag-1 autocorrelation of pre-collapse and collapse dominant species) based on their performances. We distinguished between small-scale signals (sharp drops and peaks) and large-scale signals (trend changes). Small-scale signals and large-scale signals can be at the same time, however, small-scale signals can also precede large-scale signals. Small-scale signals indicate environmental events. Large-scale trend changes refer to the decline and eventually the collapse of the community. Our results show that the collapse indicators of the dominant species outperform community-level critical slowing down indicators, which suggests that the dominant species probably have an important role in community-level collapses triggered by environmental events. We also concluded that unusual environmental events might be the number one cause of collapses therefore small-scale signals should be involved in analyses.

## Introduction

It has been long debated whether the different causes of events evoke different dynamics and patterns of collapses. This study aims to find similarities among collapses. We share the views by Erwin [1] that collapses have a primary abiotic and a secondary biotic aspect. Unusual environmental events cause unfavorable environmental circumstances, which in return, evoke structural changes of ecological communities leading to a collapse. We assume that ecological communities give distinctive and detectable signs at the environmental events and the important structural changes of the communities, respectively.

Unusual environmental events are sudden and of great magnitude. They are usually pulse events during which system parameters have a sudden positive or negative change. Bolide events, volcanic eruptions, sudden cooling or warming, fires, etc. are typical pulse events. Press events that are long-term and gradual changes in system parameters (e.g. climate change) can include unusual events as well. For instance, a sudden great increase or decrease in the temperature during gradual long-term warming or cooling is also an unusual environmental event. These events probably have an important role in the initiation of ecological collapses. They are stochastic and it is difficult or impossible to predetermine their time windows. In this study, we involved the KPG bolide event; a presumed warming event before the Paleocene-Eocene Thermal maximum; a sudden cooling and a presumed warming event during the Early Miocene, respectively; and modern, Arctic warming events during the 1920s–1940s.

Some species have a greater effect on the community structure than others do. Dominant species, the most abundant species of communities, are essential in maintaining the communities [2]. It is important to note that no uniform definition for the dominant species exists in the literature. Some based simply on their high abundance (e.g. [3–5]), others also emphasize their high impact within the community (e.g. [6–8], [2]). They probably have a fundamental role in maintaining ecosystem functions and diversity. The removal of dominant species reduces ecosystem functions and affects the diversity greatly [9]. According to Grime [10], the effect of a species on an ecosystem function is proportional to its relative abundance in a community. Based on this concept, we assume that the most abundant species is the most influential species among abundant species in a community. In this study, we apply the phrase of dominant species only to the most abundant species of the communities. The decline of dominant species causes changes in the occurrence and distribution of other species, modifies the processes and the structures of communities. Their abundance probably also provides a delay in collapses after the environmental events. The most abundant species are likely to be sensitive to environmental changes, because they gain their abundance by adapting the best to their environment. Arnoldi, Loreau and Haegeman [11] suggest that common species generate the response of a community to environmental disturbances. Significant, negative environmental changes (unusual, extreme events) affect the abundance of the dominant species greatly. According to Parmesan, Root and Willig [12], extreme events provoke a wide range of ecological responses scaling from the gene to the ecosystem, and they are important components of population extinctions. We presume that collapse triggering environmental events cause a sharp drop in the abundance of the dominant species close to the event. These events, especially pulse events, may push the abundance of the dominant species under a critical level. Overshoots may follow these undershoots after the events. The decline of the dominant species because of unfavorable environment can be beneficial for the disaster species. They can appear before the collapse, which can cause an increase in the species richness. This phenomenon can be observed under recent climate change, as well. Tropical and boreal species expand their ranges and invade into temperate and polar communities, respectively [12–13]. Lowland species are moving to higher elevations[14–16]. We suppose that a sudden increase in the species richness is often related to an unusual environmental event.

Statistical indicators can describe the history of a community. They detect the general pattern and distinctive changes in the community pattern. We suggest that collapse indicators give a short, small-scale sign (a sharp drop or a peak) at the environmental events and a large-scale sign (changes in trends) as a result of the subsequent biotic crisis. These short signs may precede the large-scale trend changes of indicators, hence involving them into collapse analyses might be important. We propose that not only the critical slowing down (CSD) indicators (see e.g. [17–18]) but also more traditional indicators can give useful information on collapses. We involved PCA (principal component analysis), HCA (hierarchical clustering analysis), the relative abundance of species, abundance, species richness, dominance, constancy, Shannon’s H in addition to the CSD indicators (lag-1 autocorrelation, variance, standard deviation) into our analysis at a community level. Considering the community structuring effect of the dominant species, we believe that the indicators of the dominant species are more powerful than community-level indicators. The dominant species affect the patterns of communities, hence all the collapse indicators.

Involving a great number of indicators into collapse analyses might be important to be able to reveal the structural changes of communities and the different aspects of collapses. Authors should also consider developing or further developing collapse indicators independently of CSD indicators to uncover new aspects of collapses. We developed two indicators (the total change of the dominant species and the lag-1 autocorrelation of the dominant species) at the level of the dominant species and one indicator (the lag-1 autocorrelation of Shannon’s H) at a community level.

To detect universal signals of collapses, we included five data series (the relative abundances of species) of different data at different levels. Our results show that the indicators of dominant species outperform the community-level indicators. We believe that distinctive, small-scale warning signs indicate collapse triggering, unusual environmental events.

### Glossary

We introduce/reintroduce some phrases for community collapses in this study in our understanding.

*Community collapse* – The collapse of the dominant species, the rise of a new dominant species and the complete structural changes in the community indicate community collapse. We initiate the use of ‘community collapse’ phrase for both community shifts and extinctions

based on common features.

*Pre-collapse community* – Pre-collapse communities cover a period starting before the environmental event and ending at the collapse boundary.

*Collapse community* – Collapse communities follow pre-collapse communities and they occupy the collapse zones where the pre-collapse dominant species are not the most abundant species anymore.

*Dominant species* – Dominant species is the most abundant species of a community in this study. Its decline is an important biotic sign of a forthcoming collapse. We use pre-collapse dominant and collapse dominant species phrases in the study

*Collapse zone vs collapse point* – We used collapse zones instead of the points of the collapse (shift or extinction) because we did not intend to include collapse zone signals into the evaluation of indicators

*Collapse zone* – We used the HCA clusters and the relative abundance of dominant species to detect the collapse communities and the collapse zones (a period of a collapse).

*Small-scale collapse signals* – We use the ‘small-scale collapse signals’ phrase for short, sharp signs (sharp drop or increase) at the environmental events or right after the events (or with a short lag). Small-scale signals precede large-scale signals (right before them or with a gap), or they are at the same time as the large-scale signals. Sometimes, short-scale signals do not precede large-scale signals. Small-scale signs indicate environmental events.

*Large-scale collapse signals* – Large-scale collapse signals are usually large-scale trend changes of the indicators referring to complete structural changes of the community. Large-scale signals are the signs of collapses.

## Materials and methods

### Data

To locate collapses and shifts, and to detect warning signs, we used open paleoecological data series available on the *PANGAEA - Data Publisher for Earth & Environmental Science* open access data library (https://pangaea.de/) [19], as well as modern paleolimnological data obtained from other authors [20]. The five studied data series range from the Cretaceous- Paleogene impact event to a recent shift including different faunal records at different time scales. These data series are supplements to peer-reviewed articles which include the change points of time series (collapses) allowing us to check the accuracy of our results. All of them show abrupt, sudden changes in the community structures.

The high-resolution Cretaceous-Paleogene record of deep-sea benthic foraminifera derives from depths between 1500–2000 m at the Northwest Pacific Ocean Drilling Program (ODP) Hole 198–1210A (Shatsky Rise, 32.2235°N, 158.2593°E, Fig 1) [21]. Shatsky Rise is a large oceanic plateau 1500 km east of Japan. The sedimentological record shows a transient assemblage change of benthic foraminifera at the Cretaceous-Paleogene (K/Pg) boundary (between 219.9–219.8 mbsf ∼ 65.489 Ma before present) probably as a result of an impact event [22]. The percent relative abundances of 101 species were collected between 215.00-–223.30 mbsf (meters below seafloor) corresponding with age between 64.70–66.14 Ma and consisting of 26 time-steps (Fig 2).

**Fig 1.**
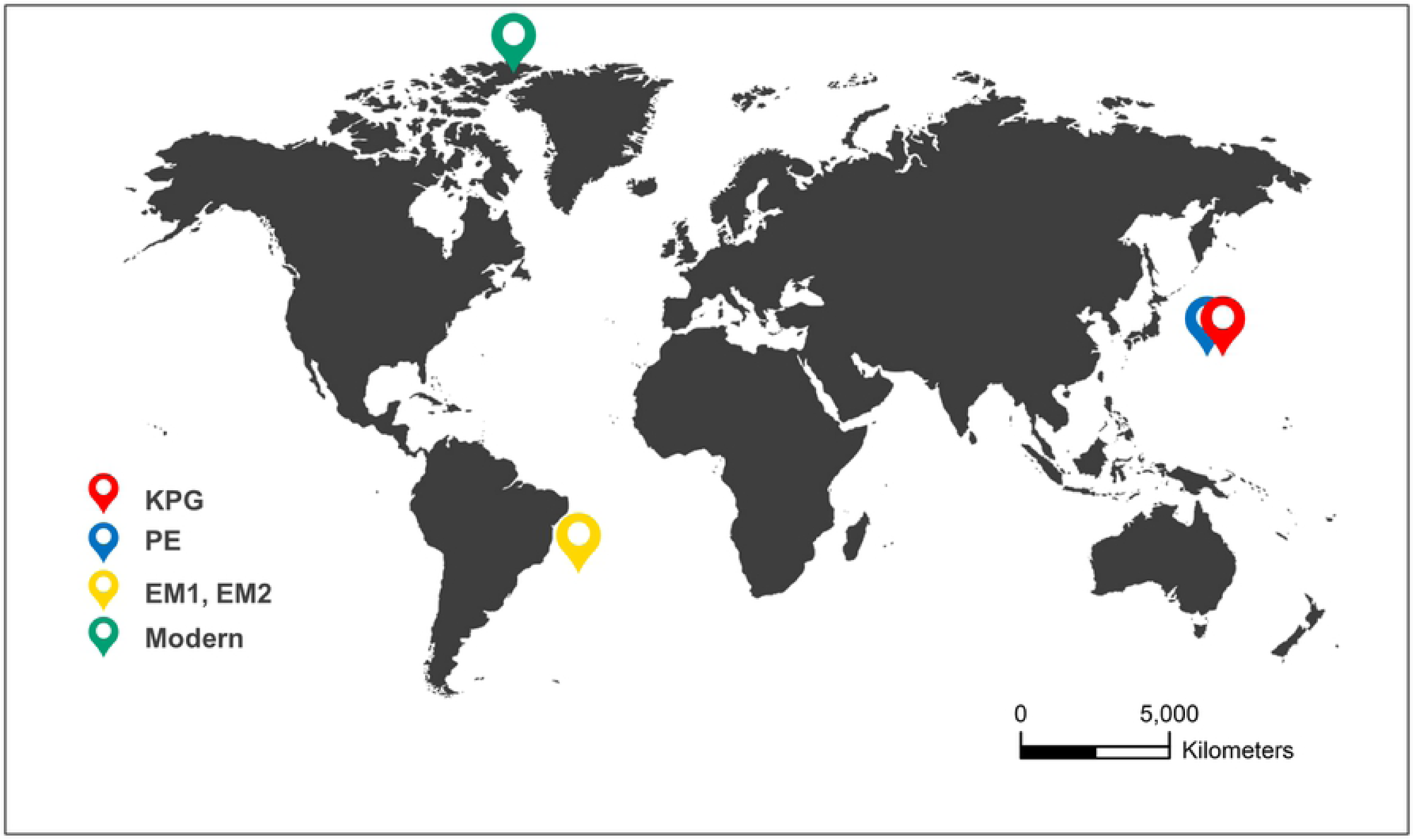
Locations of paleoecological and modern records. KPG = Cretaceous-Paleogene record of deep-sea benthic foraminifera. Shatsky Rise, northwest Pacific Ocean. Source: Alegret and Thomas [21]. PE = Paleocene-Eocene record of deep-sea benthic foraminifera. Shatsky Rise, northwest Pacific Ocean. Source: Takeda and Kaiho [23]. EM1, EM2 = Early Miocene record of deep-sea nannofossils. South Atlantic Ocean. Source: Plancq et al. [24]. Modern = Modern record of diatoms. Lake Hazen, northern Ellesmere Island, Canada. Source: Köck et al. [25], Lehnherr et al. [20]. Source of world map template: www.pptbackgroundstemplates.com.

**Fig 2.**
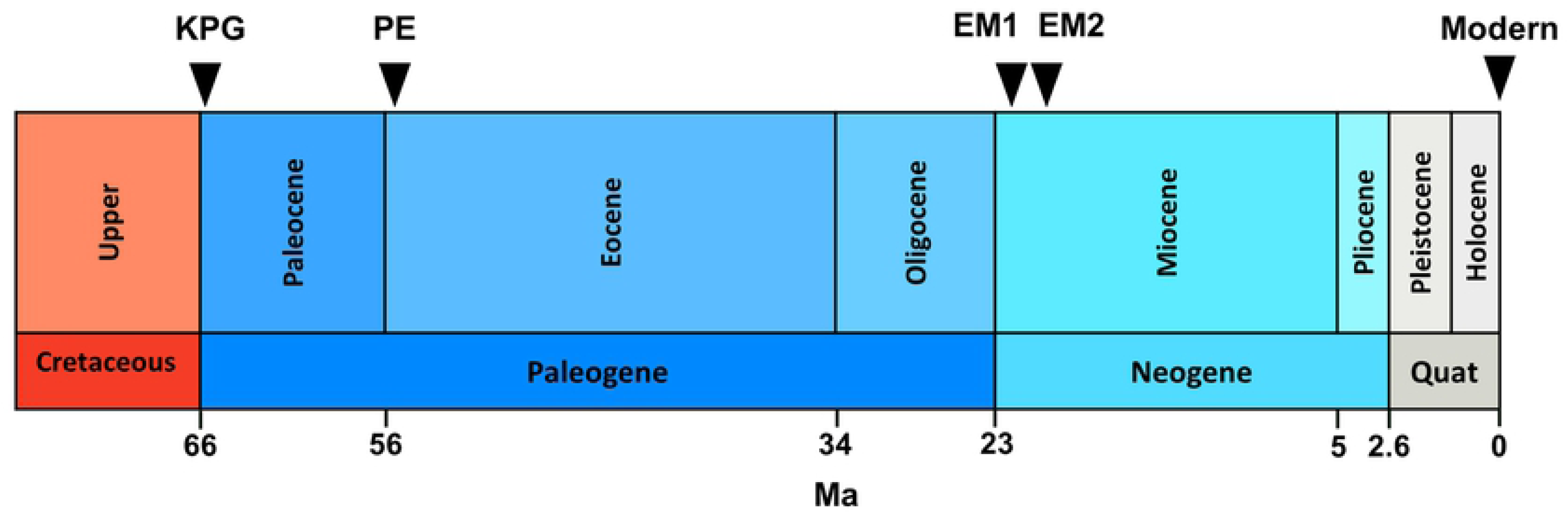
Timeline of environmental events and collapses. KPG = Extinction of benthic foraminiferal species at the Cretaceous-Paleogene boundary [21]; probable impact event (65.7 Ma). PE = Extinction of benthic foraminiferal species during the Paleocene-Eocene thermal maximum [23]; probable warming event before PETM (55.08 Ma). EM1 = Early Miocene nannofossil extinction [24]; probable glaciation event (21.3 Ma). EM2 = Early Miocene nannofossil extinction [24]; probable warming event (20.2 Ma). Modern = recent diatom shift (1995–1997) [20]; probable warming events (1924; 1939). Ma = million years ago.

**Fig 3.**
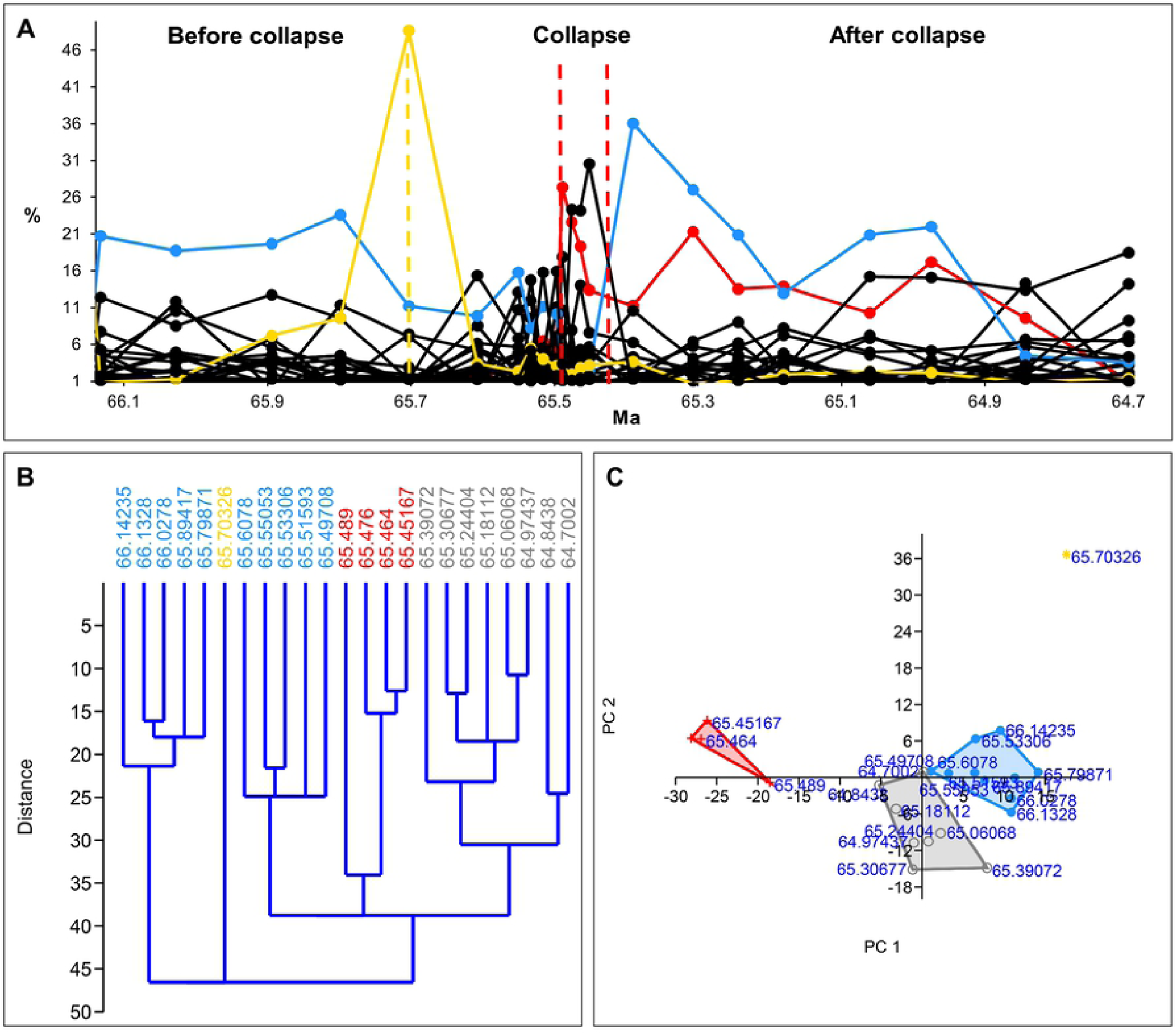
Relative abundance, hierarchical cluster analysis (HCA) and principal component analysis (PCA) of the benthic foraminiferal species across the K/Pg transition (KPG) (Northwest Pacific ODP). (A) The relative abundance of benthic foraminiferal species plot. Blue solid line = pre-collapse dominant species (*Nuttallinella ripleyensis*). Red solid line = collapse dominant species (*Bulimina kugleri)*. Yellow solid line = environmental indicator (*Pyraminida rudita*). Yellow dashed vertical line = environmental event (bolide event). Red dashed vertical line = collapse zone boundary. (B) HCA plot and (C) PCA plot. Blue = pre- collapse community, red = collapse community, grey = recovery community, yellow = environmental event (bolide event). Source of data: Alegret and Thomas [21].

The Paleocene-Eocene sediments were collected from ODP Hole 1212B (Shatsky Rise, 32.4485°N, 157.7117°E, water depth 2681 m, Fig 1) [23]. The high-resolution analysis revealed a major benthic foraminiferal extinction event during the Paleocene-Eocene thermal maximum (PETM) [26]. The PETM interval at ODP Hole 1212B was between 55.0 Ma–54.9 Ma (79.925–79.700 mbsf) [23]. The relative abundances of 27 species come from a depth between 79.705–80.005 mbsf covering a period between 54.90–55.21 Ma with 31 time-steps (Fig 2).

The late Oligocene-early Miocene deep-sea nannofossil data come from the South Atlantic Deep Sea Drilling Program (DSDP) Site 516 (30.2763°S, 35.2852°W, water depth 1313 m, Fig 1) situated about 1000 km away from the Brazilian coasts in the South Atlantic Ocean [24].

The relatively warm climate of Early Miocene was interrupted with transient Antarctic glaciation events (Mi-1a and Mi-1aa events at about 21.3 Ma /approx. 170–180 mbsf/ and 20.2 Ma /approx. 150–160 mbsf/ respectively), which may be shown by the decline of *Cyclicargolithus floridanus*, the dominant species, in the record [27]. The nannofossil assemblage was sampled between 104.46–239.05 mbsf which covers the period between the Oligocene-Miocene boundary and the early Miocene (Fig 2). It includes the relative abundances of 13 species with 33 time-steps. We used two periods of the data series for analysis between 161.48–203.2 mbsf (EM1) and 109.78–157.45 mbsf (EM2), respectively. Both of them belong to the Early Miocene (EM) sub-epoch. EM1 probably includes a glaciation event and EM2 involves a warming event.

The modern data series originates from a paleolimnological record of diatoms. The paleolimnological record of diatoms comes from the microfossil analysis of three sediment cores collected from Lake Hazen in 2013 (northern Ellesmere Island, Canada, near the deepest point of the lake /81.816806° N, 70.700222° W, Fig 1/, water depth 260 m) [25]. Lake Hazen is the largest lake north of the Arctic Circle situated in Quttinirpaaq National Park. The recent climate warming resulted in an ecological shift in the lake. The algal (diatom) primary producers shifted from shoreline benthic to open-water planktonic species probably because of the earlier onset of ice melting after winters and the longer growing seasons with a larger ice-free area between 1995–1997 [20]. The warming of the Lake Hazen watershed started more intensively around the turn of the twentieth century. The period between 2000 and 2012 was especially warm with a 2.6 °C increase in the mean summer land surface temperatures of ice-covered regions [20]. The relative abundances of 63 species refer to the period of 1894– 2014 (top 10–20 cm of the cores) with 18 time-steps (Fig 2).

## Methods

To locate the collapses in the data series and to define the possible warning signals, we applied both univariate and multivariate indicators at the community level and the level of the dominant species.

### Community and unusual event indicators

We applied the relative abundance of species (%), HCA (hierarchical cluster analysis) and PCA (principal component analysis) to define the communities and unusual environmental events. We used multidimensional methods, temporal clustering and principal component analysis (PCA), for dimension reduction in Past v 2.17 [28]. First, we applied time series clustering to detect communities and unusual events (environmental perturbations, extreme events). In our hierarchical cluster analyses (HCA), we used an unweighted pair-group average (UPGMA) algorithm, for which the distance matrix was computed using Euclidean distance. We preferred constrained clustering so that the adjacent rows of time series are joined during the agglomerative clustering. The products of our cluster analysis are dendrograms showing the clustered data points which represent the communities (before collapses, during collapses, after collapses) in the time series. We presume that the dendrograms also represent unusual environmental events as outliers.

We used principal component analysis (PCA) to explore the similarities between clusters (communities). PCA shows how the communities are related to each other and they provide information about the environmental events (climate change, unusual environmental events). PCA was done by eigenvalue decomposition of a data covariance matrix. The convex hulls in the PCA scatterplots represent the communities.

The relative abundance of species (%) shows the unusual environmental events as a peak of the environmental indicator species and as a sharp drop of the pre-collapse dominant species at the same time. The boundary between the pre-collapse and the collapse communities is at the change point where the pre-collapse dominant species loses its absolute dominance (not the most abundant species any more).

### Collapse indicators at the community level

At the community level, we applied the following collapse indicators: *total abundance, species richness, dominance, constancy, Shannon’s H*, as well as the indicators of critical slowing down (*standard deviation, variance, lag-1 autocorrelation)*. *Total abundance* refers to the number of individuals in a community. *Species richness* is the number of species in a community. *Dominance* is the degree to which the dominant species is more numerous than other species of the community. It reflects the effect of the dominant species on the community.

A formula for determining the degree of dominance:

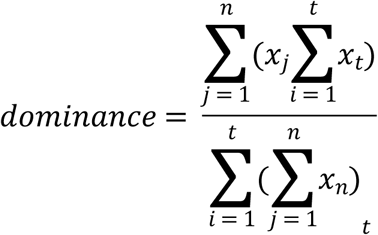

where

x = relative abundance data of species (%) n = number of species

t= number of time steps

*Constancy* is the dispersion of species in a community. It also reflects the effect of the dominant species on the community.

A formula for determining constancy:

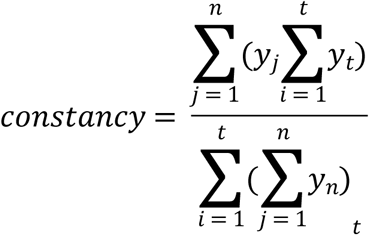

where

y = presence/absence data of species

n = number of species

t= number of time steps

The temporal changes in *Shannon’s H* [29] provides information on diversity and community composition changes. The index is based on the abundance and the evenness of the species. Shannon diversity index mostly ranges from 1.5 to 3.5. The greater the number, the higher diversity the community has.

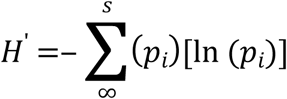

*Standard deviation, variance* and *lag-1 autocorrelation* are the indicators of critical slowing down, e.g. [17–18]. ‘Critical slowing down’ (CSD) is a phenomenon of dynamical systems near critical transitions. It was first described in the catastrophe theory in the 1970s (e.g. [30]) and it was reintroduced in ecology in the last decades. At critical thresholds or tipping points (bifurcation), the complex dynamical systems shift abruptly from one state to another, which we call critical transition. Critical transitions or abrupt shifts can be observed in complex systems from medicine [31], through finance [33–34] to the climate system [35–36]. During mass extinctions and ecological shifts, the systems undergo a critical transition, as well. Once the system gets close to the critical threshold or tipping point, it slows down, which means that it takes the system a longer time to recover from small disturbances. The cause of this slowness in response is the low level of resilience. Commonly used CSD indicators are variance, lag-1 autocorrelation and skewness. According to Dakos *et al*. [36], CSD causes a rise in variance and temporal correlation. Flickering may be a reason for increasing variance [38–39]. Flickering is the quick alternating of a system between two different states before the critical transition.

In the past few years, authors have become more critical with CSD indicators. CSD indicators do not always increase before critical transitions [37,39]. Guttal, Jayaprakash and Tabbaa [40] suggest that variance is not a reliable CSD indicator, because it either decreases or increases before regime shifts. Sometimes, regime shifts have no tipping points and in this case, there is no CSD at regime shifts [37]. Many range shifts are long and smooth after the tipping point and it is difficult to detect them, which delays management actions [41]. Burthe *et al*. [42] tested CSD indicators on 125 long-term abundance time-series of 55 taxa “across multiple trophic levels in marine and freshwater ecosystems” and they concluded that they are not effective in predicting nonlinear changes. Biggs, Carpenter and Brock [43] and Spanbauer *et al*. [44] propose that a large increase in collapse indicators happen at the onset of collapses, which is too late for intervention.

We applied both standard deviation and variance to measure the temporal variance in data series prior to collapses. Both of them are the indicators of critical slowing down, e.g. [17–18]. We intend to test the effectiveness of these two indicators and to make a comparison between them.

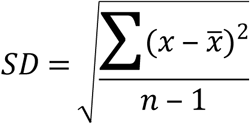

We expressed variance as *CV* = *SD*/*mean* [45].

We applied the Pearson correlation [46] as a measure of lag-1 autocorrelation between subsequent time steps. Pearson correlation is considered to be effective as an indicator of critical slowing down [17]. Having time series with relatively few time steps (18–33 time steps), we applied a moving window size 3.

We introduce a new collapse indicator at the community level, the lag-1 autocorrelation of Shannon’s H. We hypothesize that lag-1 autocorrelation of Shannon diversity index is sensitive to changes leading to collapses, therefore it is suitable for detecting the beginning of tendentious structural changes of communities. We presume that the community composition and the evenness of species change greatly at the environmental events triggering a collapse, which can be measured with Shannon’s H index. We also suspect that the correlation between the subsequent time steps changes significantly at the environmental events, therefore the lag- 1 autocorrelation of subsequent time steps probably also changes largely. Hence, we expect a strong warning signal of the lag-1 autocorrelation of the Shannon diversity index.

The lag-1 autocorrelation of Shannon’s H varies between -1 and +1. The formula for lag-1 autocorrelation of Shannon’s H is based on the formula of Pearson correlation:

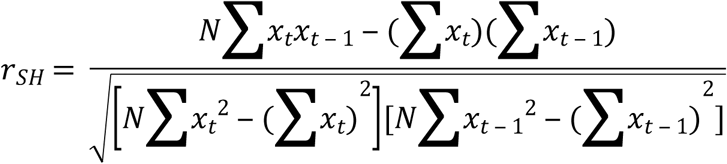

where

x_t_ = Shannon diversity index at time step ‘t’

x_t-1_ = Shannon diversity index at time step ‘t-1’

### Collapse indicators at the level of the dominant species

We presume that collapse indicators at the level of the dominant species are more effective in detecting the warning signs of collapses than indicators at the community level because the dominant species are probably sensitive to collapse triggering environmental events and they are one of the most influential species in a community (see *Introduction*). We introduce two collapse indicators at the level of the dominant species: the *total change of pre-collapse and collapse dominant species (TCD)* and the *lag-1 autocorrelation of pre-collapse and collapse dominant species*.

*The total change of dominant species* (*TCD*) shows the degree of change in the abundance of dominant species relative to the first time step (presumably before the environmental events) over time.

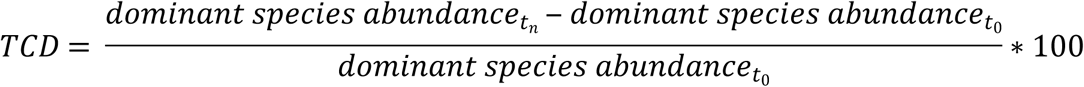

where

t_0_ = the first time step of data series

t_n_ = time step

We presume that the sharp drop of the total change of the dominant species is a sign of an upcoming collapse.

*Combined indicator of the total change of pre-collapse and collapse dominant species*. We created a combined indicator of the total change of dominant species by considering always the earlier warning sign of either pre-collapse or collapse species (see *Evaluation of collapse indicators*).

We measured the *lag-1 autocorrelation of pre-collapse and collapse dominant species* applying Pearson correlation to test the lag-1 autocorrelation CSD indicator at the level of the dominant species. This indicator measures the correlation between the subsequent time steps of the dominant species abundance.

*Combined indicator of lag-1 autocorrelation of pre-collapse and collapse dominant species*. We created a combined indicator of lag-1 autocorrelation of dominant species by considering always the earlier warning sign of either pre-collapse or collapse species *(see Evaluation of collapse indicators*).

We applied Excel 2016, R 3.6.2 and ArcMap 10.4 for data visualization. We used R software [47] to create the figures of the collapse indicators (see *Evaluation of collapse indicators*). We applied *ggplot2* and *reshape2* packages for the faceted plots. To create the R Markdown files for the faceted plots, we used the *rmarkdown* package. We georeferenced the world map of Fig 1 in ArcMap 10.4.

## Results

### Communities and boundaries

We applied the HCA and the relative abundance of species to identify the clusters and the boundaries between them. The clusters/sub-clusters represent the communities.

#### KPG

In the KPG data series, the pre-collapse community is in the interval between 66.14235– 65.49708 Ma. This period includes two clusters between 66.14235–65.79871 Ma (cluster 1) and 65.6078–65.49708 Ma (cluster 2), respectively, as well as the bolide event between them 65.70326 Ma. Looking at the figure of the ‘Relative abundance of the benthic foraminiferal species across the K/Pg transition’ and the HCA (Figs 3A and 3B), cluster 2 gives the expression of a collapse zone. However, cluster 1 and cluster 2 have the same dominant species, namely *Nuttallinella ripleyensis* [21], hence we suppose that cluster 2 belongs to the pre-collapse community. The collapse zone is cluster 3 in which the dominant species is *Bulimina kugleri* [21].

#### PE

The HCA clearly indicates the pre-collapse community (80.005–79.915 mbsf) and the collapse community (79.905–79.855 mbsf) (Fig 4B). The pre-collapse dominant species is *Siphogenerinoides brevispinosa* [23]; the collapse dominant species is *Bolivina gracilis* [23].

**Fig 4.**
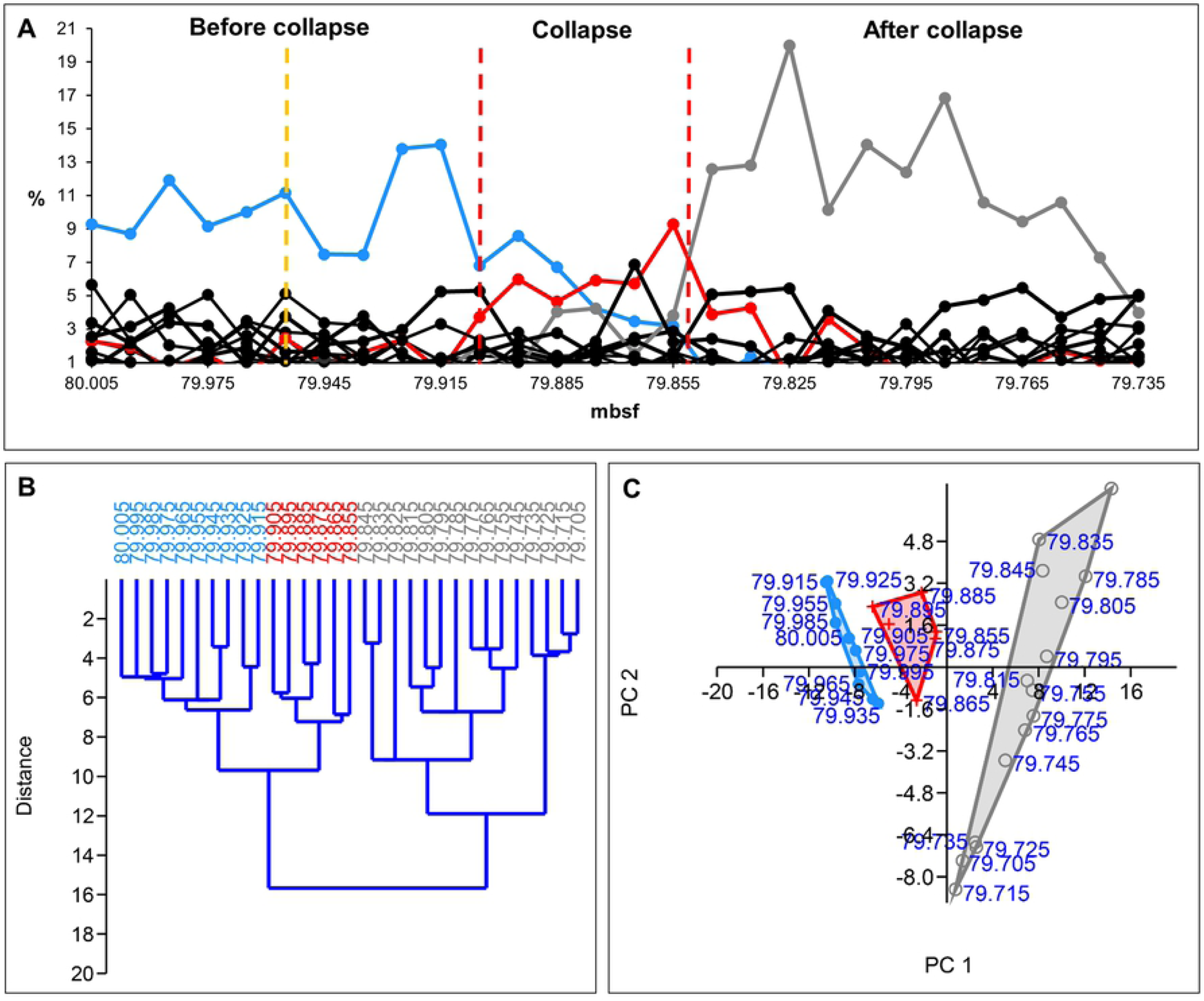
Relative abundance, hierarchical cluster analysis (HCA) and principal component analysis (PCA) of Paleocene-Eocene benthic foraminifera (Central Pacific, ODP Hole 1212B). (A) The relative abundance of benthic foraminiferal species plot. Blue solid line = pre-collapse dominant species (*Siphogenerinoides brevispinosa*). Red solid line = collapse dominant species (*Bolivina gracilis*). Grey solid line = recovery dominant species (*Quadrimorphina profunda*). Yellow dashed vertical line = presumed environmental event (temporary warming). Red dashed vertical line = collapse zone boundary. (B) HCA plot and (C) PCA plot. Blue = pre-collapse community, red = collapse community, grey = recovery community. Source of data: Takeda and Kaiho [23].

#### EM1 and EM2

The HCA of the original Late Oligocene-early Miocene data series shows four communities (community 1–4) (Fig 5B). EM1 data series includes the collapse of community 1 (pre-collapse community) (Fig 6). The HCA shows community 1 as cluster 1 between 203.2–171.93 mbsf. However, the relative abundance of species suggests that the boundary between community 1 and community 2 is earlier, between 177.92–172.79 mbsf. After 177.92 mbsf, *Dictyococcites spp*. [24], the dominant species of community 2 take over the dominance from *Cyclicargolithus floridanus* [24], the dominant species of community 1. Hence, we decided to modify the boundary between community 1 and community 2 suggested by the HCA. EM2 involves the collapse of community 3 (pre-collapse community) (Fig 7). The HCA shows community 3 as cluster 3 between 157.45–137.21 mbsf. The pre-collapse dominant species of EM2 is *Dictyococcites antarcticus* [24]. The collapse dominant species are *Coccolithus spp*. [24].

**Fig 5.**
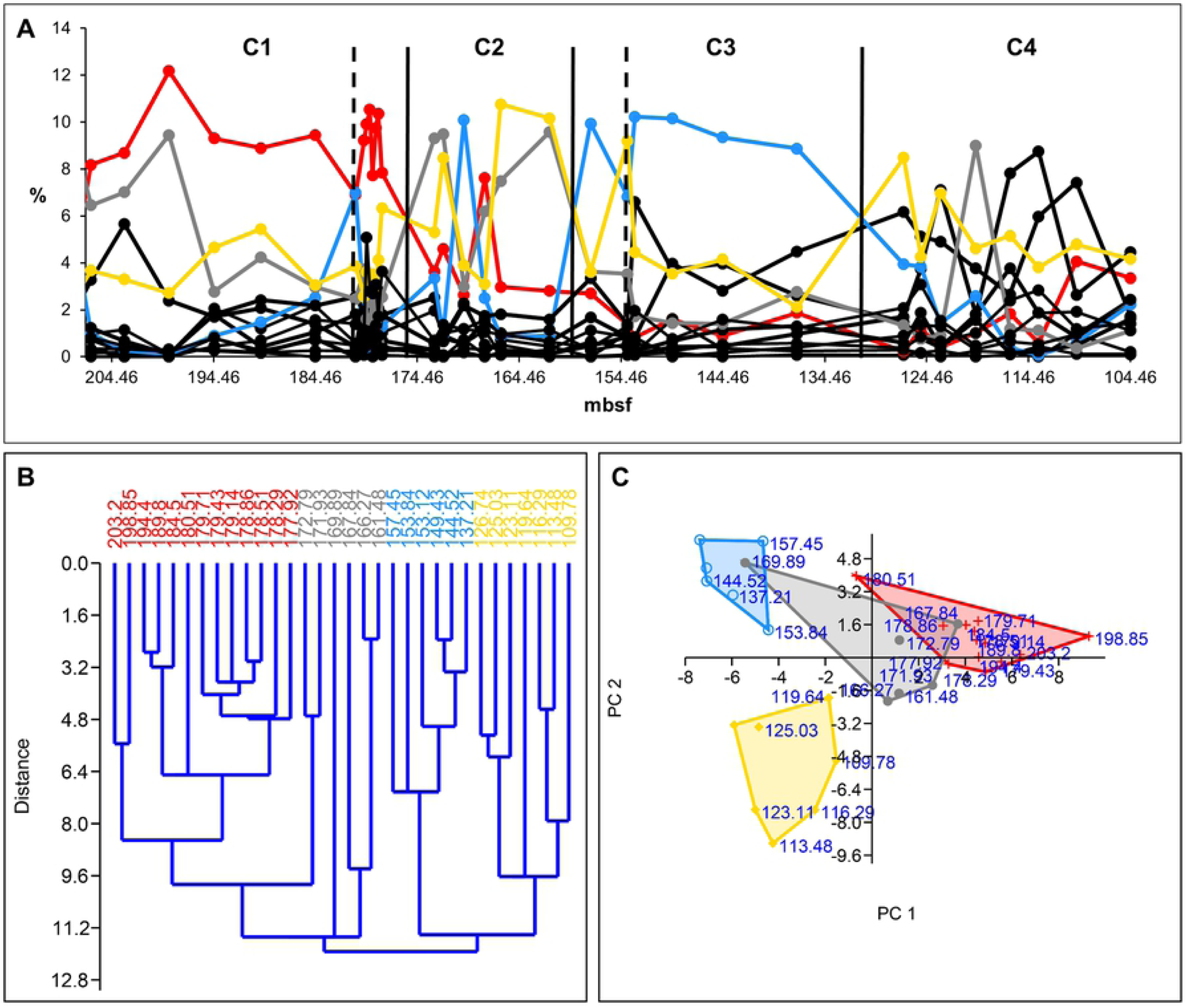
Relative abundance, hierarchical cluster analysis (HCA) and principal component analysis (PCA) of late Oligocene – Early Miocene nannofossils (South Atlantic DSDP Site 516). (A) The relative abundance of nannofossils. C1 = community 1 (EM1 pre-collapse community), C2 = community 2 (EM1 collapse community), C3 = community 3 (EM2 pre-collapse community), C4 = community 4 (EM2 collapse community). Red solid line = dominant species 1 (*Cyclicargolithus floridanus*). Grey solid line = dominant species 2 (*Dictyococcites spp*.). Blue solid line = dominant species 3 (*Dictyococcites antarcticus*). Yellow solid line = dominant species 4 (*Coccolithus spp*.). Black solid vertical line = community boundary, black dashed vertical line = environmental event. (B) HCA plot and (C) PCA plot. Red = community 1, grey = community 2, blue = community 3, yellow = community 4. Source of data: Plancq et al. [24].

**Fig 6.**
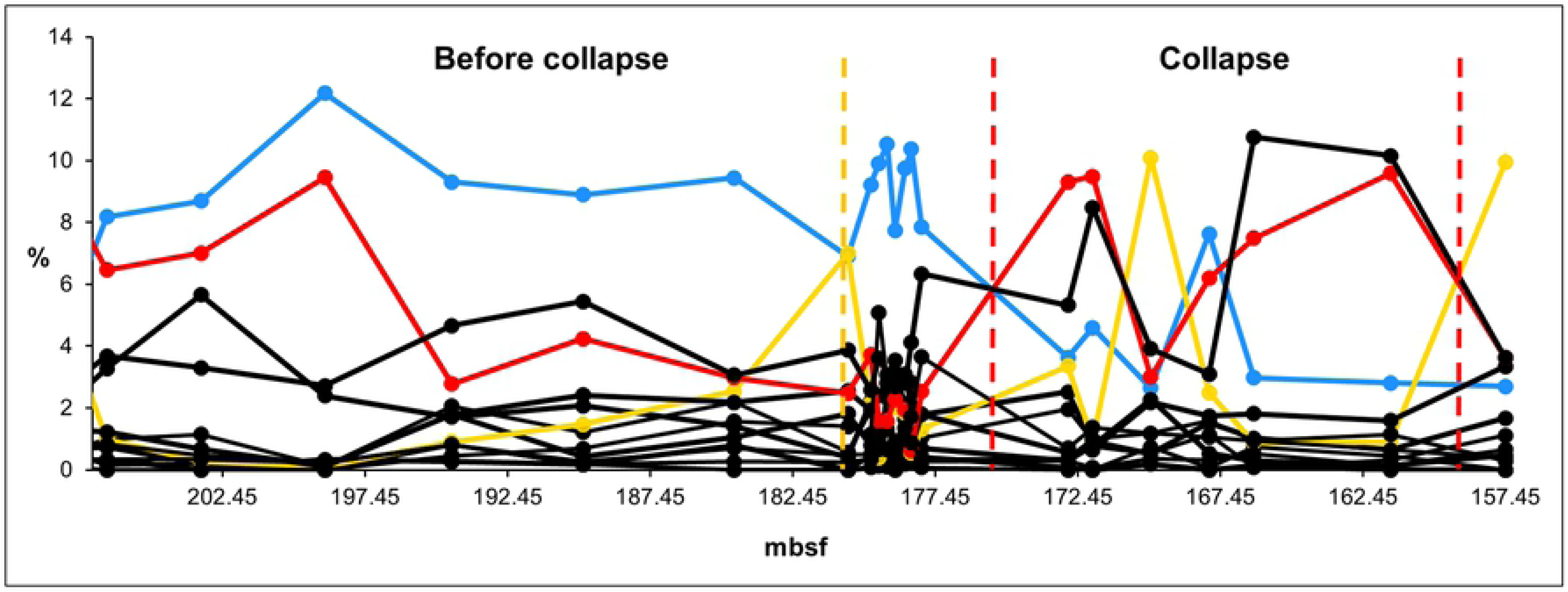
Relative abundance of Early Miocene nannofossils (EM1) (South Atlantic DSDP Site 516). Blue solid line = Pre-collapse dominant species (*Cyclicargolithus floridanus*). Red solid line = Collapse dominant species (*Dictyococcites spp*.). Yellow solid line = environmental indicator (*Dictyococcites antarcticus*). Yellow dashed vertical line = presumed environmental event (temporary glaciation). Red dashed vertical line = collapse zone boundary. Source of data: Plancq et al. [24].

**Fig 7.**
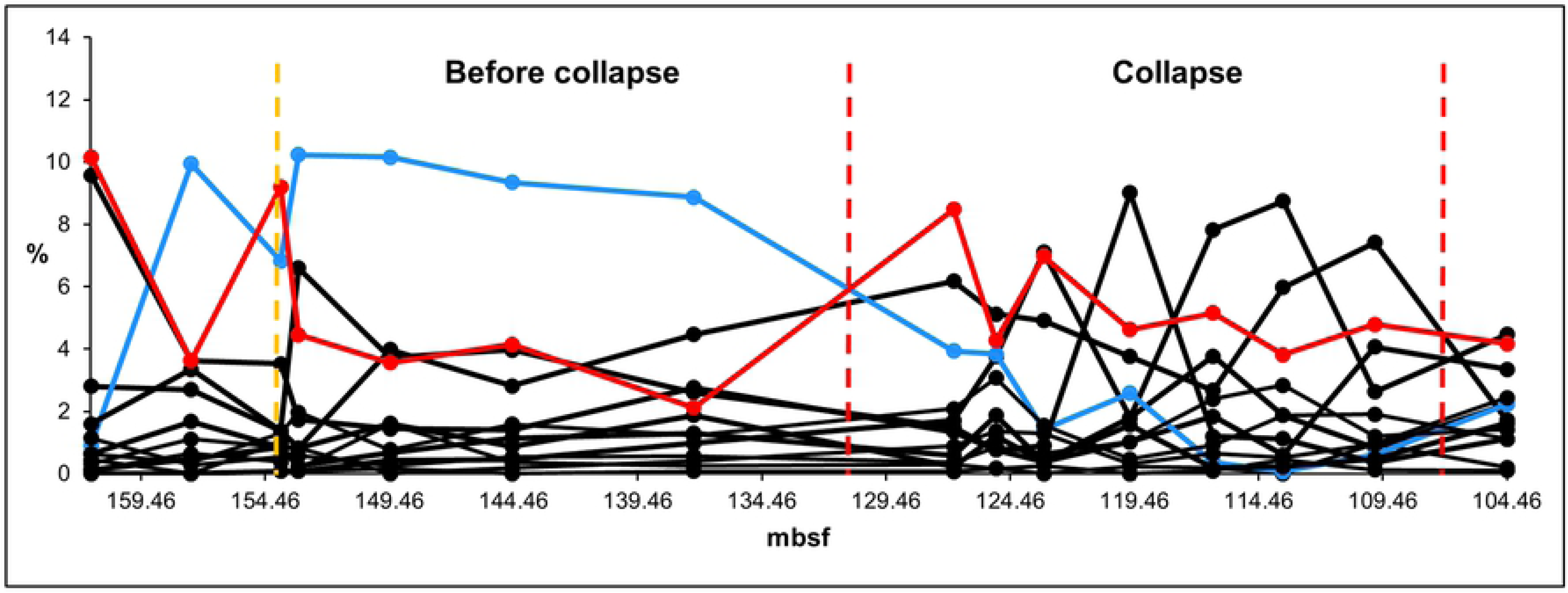
Relative abundance of Early Miocene nannofossils (EM2) (South Atlantic DSDP Site 516). Blue solid line = Pre-collapse dominant species (*Dictyococcites antarcticus*). Red solid line = collapse dominant species (*Coccolithus spp*.). Yellow dashed vertical line = presumed environmental event (temporary warming). Red dashed vertical line = collapse zone boundary. Source of data: Plancq et al. [24].

#### Modern

The modern data series shows a recent diatom shift under global warming in Lake Hazen (northern Ellesmere Island, Canada) (Fig 8). The pre-collapse community represents the period between 1894–1983 as the HCA suggests. The collapse zone is between 1991–2003. The pre-collapse dominant species is *Staurosirella pinnata* [20], the collapse dominant species is *Cyclotella comensis* [20]. The second most abundant pre-collapse species is *Staurosira construens* [20]. The ratio of *S*. *pinnata* and *S*. *construens* provides information on summer air temperature [48] (see *Identifying environmental events*).

**Fig 8.**
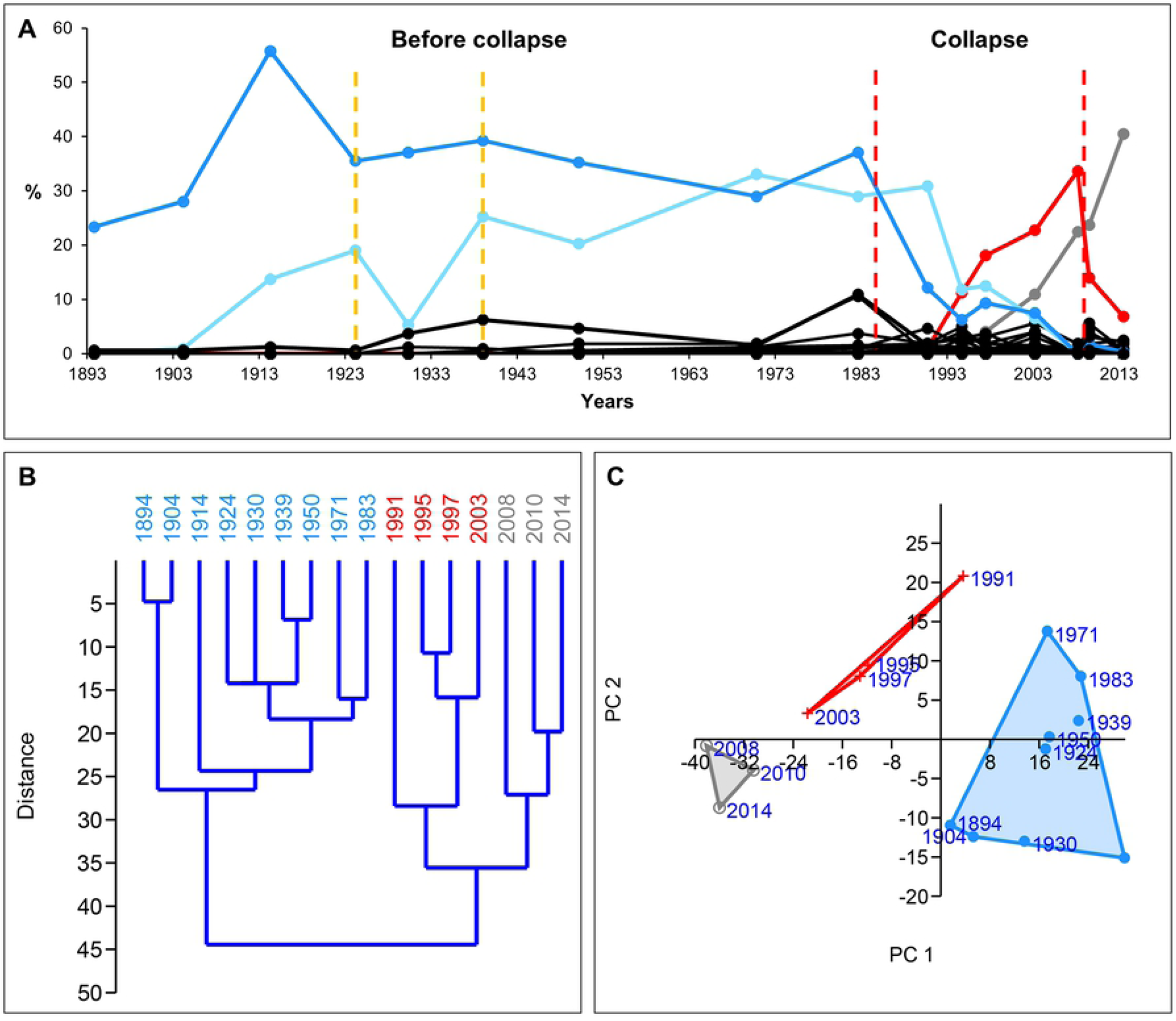
Relative abundance, hierarchical cluster analysis (HCA) and principal component analysis (PCA) of diatoms (Lake Hazen, northern Ellesmere Island, Canada). (A) The relative abundance of diatoms. Blue solid line = Pre-collapse dominant species (*Staurosirella pinnata*). Red solid line = collapse dominant species (*Cyclotella comensis*). Light blue solid line = *Staurosira construens*. Grey solid line = recovery dominant species (*Discostella stelligera*). Yellow dashed vertical line = environmental event (summer air temperature rise). Red dashed vertical line = collapse zone boundary. (B) HCA plot and (C) PCA plot. Blue = pre-collapse community, red = collapse community, grey = recovery community. Source of data: Köck et al. [25], Lehnherr et al. [20].

### Identifying environmental events

We identified the environmental events which probably contribute to the collapses using the literature, PCA, HCA, and the relative abundance of species. PCA and HCA have outliers at the events. The relative abundance of species gives small-scale, sharp signs, drops or peaks, close to the events. The relative abundance of dominant species usually drops sharply close to the environmental events. Environmental indicators usually have a peak. For instance, in the KPG data series, the relative abundance of *Pyramidina rudita* [21], an environmental indicator, has a huge peak, at the bolide event (Fig 3A). In the EM1 data series, the relative abundance of *D*. *antarcticus*, the indicator of the cooling [49], also has a high peak at the temporary glaciation event (Fig 6A).

We had two types of collapse triggering environmental events in the data series: sudden, temporary pulse events (KPG, PE, EM1, EM2); and sudden shifts in an environmental parameter, namely temperature (PE, modern).

#### KPG

The indicators of the KPG data series clearly show the time of the bolide event. The relative abundance of *P*. *rudita*, which is an environmental indicator, has a high peak 65.70326 Ma (Fig 3A). At the same time, *N*. *ripleyensis*, which is the pre-collapse dominant species, drops sharply. In the HCA, the outlier 65.70326 Ma separates the clusters before and after the bolide event (Fig 3B). In the PCA, the outlier is in the right-hand upper corner away from the clusters (Fig 3C). Time step 65.70326 Ma is an outlier different from any other time step in the data series.

#### PE

The Paleocene-Eocene Thermal Maximum was a period at around the boundary between the Paleocene and Eocene geological epochs when the temperature rose by 5–8 °C [50]. The study by Takeda and Kaiho [26] indicates the start of this environmental event at 79.925– 79.915 mbsf. At this time, *S*. *brevispinosa*, the dominant species of the pre-collapse community, shoots up. The HCA shows this period as part of cluster 1 (pre-collapse community), but as a separate sub-cluster (Fig 4B). In the HCA, 79.925–79.915 mbsf are outliers. However, based on the relative abundance of species and the HCA (Figs 4A and 4B), we identified a previous environmental event, maybe a pulse event that preceded the PETM and it may have contributed to the collapse. The HCA shows an outlier in cluster 1 at 79.955 mbsf. After this event, at 79.945 mbsf, the pre-collapse dominant species (*S*. *brevispinosa*) drops sharply, which is a general sign of environmental events affecting communities dramatically. In PCA, 79.925–79.915 mbsf and 79.955 mbsf are close to each other, which suggests a connection between these time steps (Fig 4C). A smaller environmental event might have also occurred at 79.995 mbsf as it is shown by HCA, however, it is a weak sign, and therefore we did not involve it in the analysis. We hypothesize that the event at 79.955 mbsf is a pulse event initiating/contributing to the collapse before the PETM.

#### EM1 and EM2

The HCA, the relative abundance of the Late Oligocene-early Miocene nannofossils (Figs 5A and 5B) and the total change of the dominant species (Fig 9) help to identify the environmental events and the direction of the environmental changes. During the Early Miocene (23–16 Ma), the climate started to cool. Sudden, temporary glaciation events occurred between about 21.3 Ma–20.2 Ma (between approx. 170–180 mbsf and 150–160 mbsf) [27]. The HCA shows the cooling between clusters 1–3 (Fig 5B). Cluster 4 is different from clusters 1–3. It is probably a warmer period. The total change of the dominant species of community 1–4 indicates the direction of the environmental changes and the environmental events (Fig 9). *D*. *antarcticus* indicates the temporary glaciation events by huge peaks at 180.51 mbsf (community 1) and 169.89 mbsf (community 2), respectively, as well as the glaciation period between 157.45–137.21 mbsf (community 3) (Fig 9).

**Fig 9.**
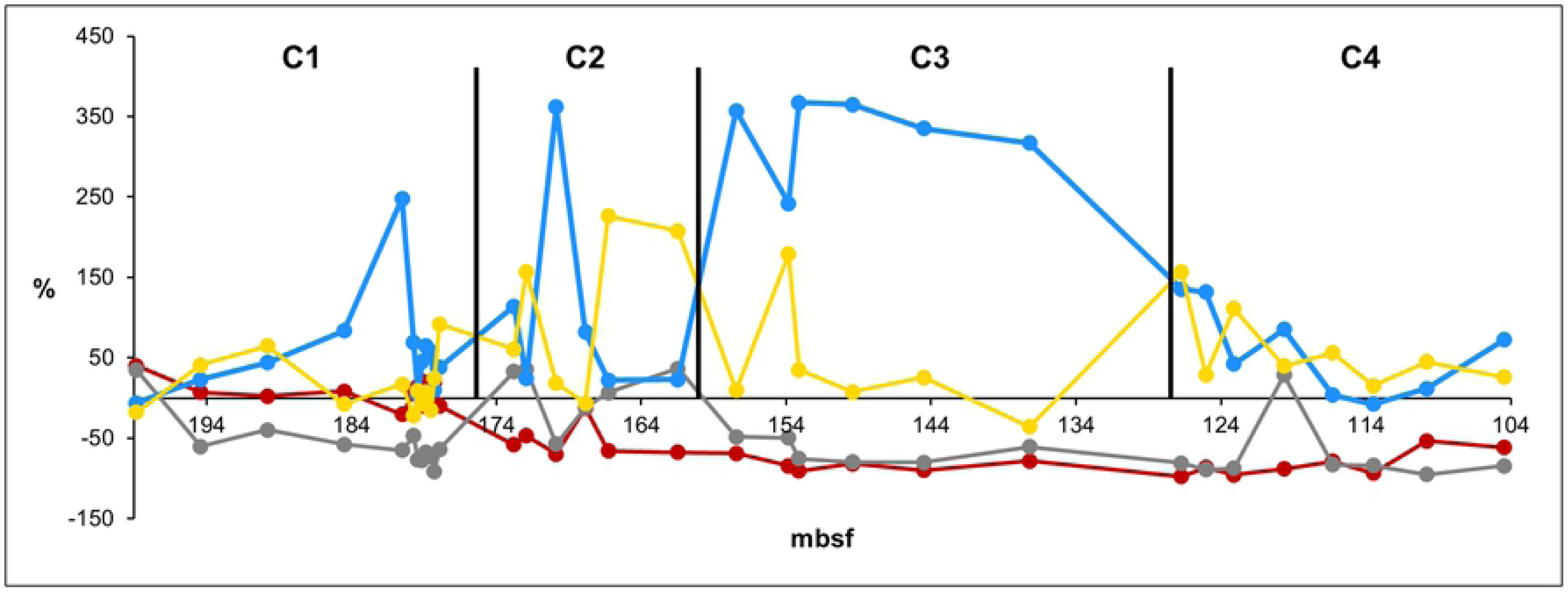
The total changes of dominant species (%) of community 1–4. C1–4 = community 1–4. Red solid line = dominant species 1 (*Cyclicargolithus floridanus*), grey solid line = dominant species 2 (*Dictyococcites spp*.), blue solid line = dominant species 3 (*Dictyococcites antarcticus*), yellow solid line = dominant species 4 (*Coccolithus spp*.). Black solid vertical line = community boundary.

EM1 data series covers a period at the beginning of Early Miocene when the climate started to cool. It includes a temporary glaciation event at 180.51 mbsf (Fig 6). EM2 data series covers a glaciation period in the Early Miocene (Fig 7). It probably has a temporary warming event at 153.84 mbsf which may have initiated/contributed to the collapse of the dominant species (*D*. *antarcticus*). At 153.84 mbsf, *D*. *antarcticus* drops suddenly, while *Coccolithus spp*. have a high peak (Fig 7). *Coccolithus spp*. are indicators of interglacial periods [51].

#### Modern

The main reason for the recent diatom shift in Lake Hazen is the warming climate [20]. Unfortunately, the study by Lehnherr et al. [20] does not include information on the local temperature data of the Hazen Lake region for the whole diatom data series, therefore we applied an indirect method to be able to detect environmental events. The increasing summer air temperature, the expansion of the growing season and the earlier melting of the ice sheet [20] are probably important contributors to Arctic melting. The reduced summer albedo due to sea ice and snow cover loss may also be a key component of Arctic amplification [52]. According to Finkelstein and Gajewski [48], the lower ratios of *Staurosirella pinnata* to *Staurosira construens* refers to warmer summer air temperatures, while greater ratios suggest cooler summers. Using this information, we can learn about the local trends of temperature and perhaps unusual environmental events. To make it more expressive, we use the reverse relationship between these two species, which means that the higher ratios of *Staurosira construens* to *Staurosirella pinnata* refer to warmer summers. In Fig 10, you can see the ratios of *Staurosira construens* to *Staurosirella pinnata* between 1904–1991. The ratio started to increase in 1904 and it shows a general increasing trend. 1924 might be an important time step because the ratio of *Staurosira construens* to *Staurosirella pinnata* passes 0.5. The ratio drops temporarily in 1930, then it stays high. After 1983, it shoots up. The local air temperature trends may also reflect regional Arctic trends. The near-surface temperature increased in the Arctic region significantly between the 1920s–1940s. According to Yamanouchi [53], Arctic warming from the 1920s to the 1940s is similar to the recent 30–40-year warming.

**Fig 10.**
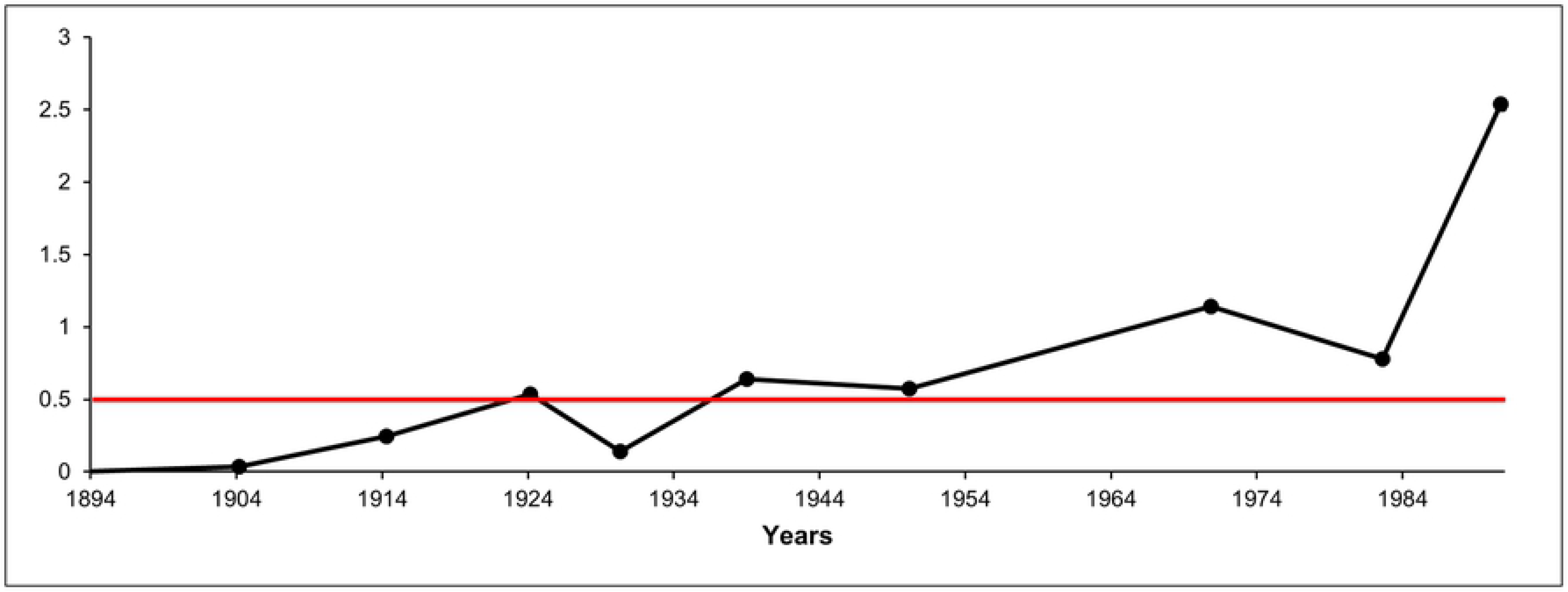
Ratios of *Staurosia construens* and *Staurosirella pinnata*, based on Finkelstein and Gajewski [48]. The increasing ratios refer to warmer summers. Red solid line = critical threshold. The ratio first rose above the threshold in 1924, and it stayed above it after 1939.

The probable temporary cooling in 1930 may be a local event (Fig 10). As the figure shows, the ratio has remained high from 1939 and it has been increasing exponentially from 1983 which marks the start of the collapse zone. We consider both 1924 and 1939 important time steps and environmental events in local warming. In 1924, the ratio of *Staurosira construens* to *Staurosirella pinnata* passes 0.5, which is a critical threshold and refers to significantly warmer summers. 1939 indicates the onset of the long-term high summer air temperatures.

### Evaluation of collapse indicators

The evaluation of the collapse indicators is between the probable environmental event and the start of the collapse zone. The modern data series includes the absence data of the collapse dominant species in the pre-collapse zone, hence the indicators of the collapse dominant species underperform.

The evaluated collapse indicators at the level of the community are as follows: *abundance, species richness, constancy, dominance, Shannon’s H, standard deviation, variance, lag-1 autocorrelation of community, lag-1 autocorrelation of Shannon’s H* (Figs. 11–15). The evaluated collapse indicators at the level of the dominant species are *the total changes of pre-collapse and collapse dominant species, respectively, and the lag-1 autocorrelation of pre-collapse and collapse dominant species, respectively* (Figs. 11–15). (Also, see *Methods* for more information on the indicators.) Own-developed indicators are the *lag-1 autocorrelation of Shannon’s H, the total change of dominant species, and the lag-1 autocorrelation of the dominant species* (Figs. 11–15). We used R software [47] to create the figures of collapse indicators. We applied ggplot2 and reshape2 packages for the faceted plots.

**Fig 11.**
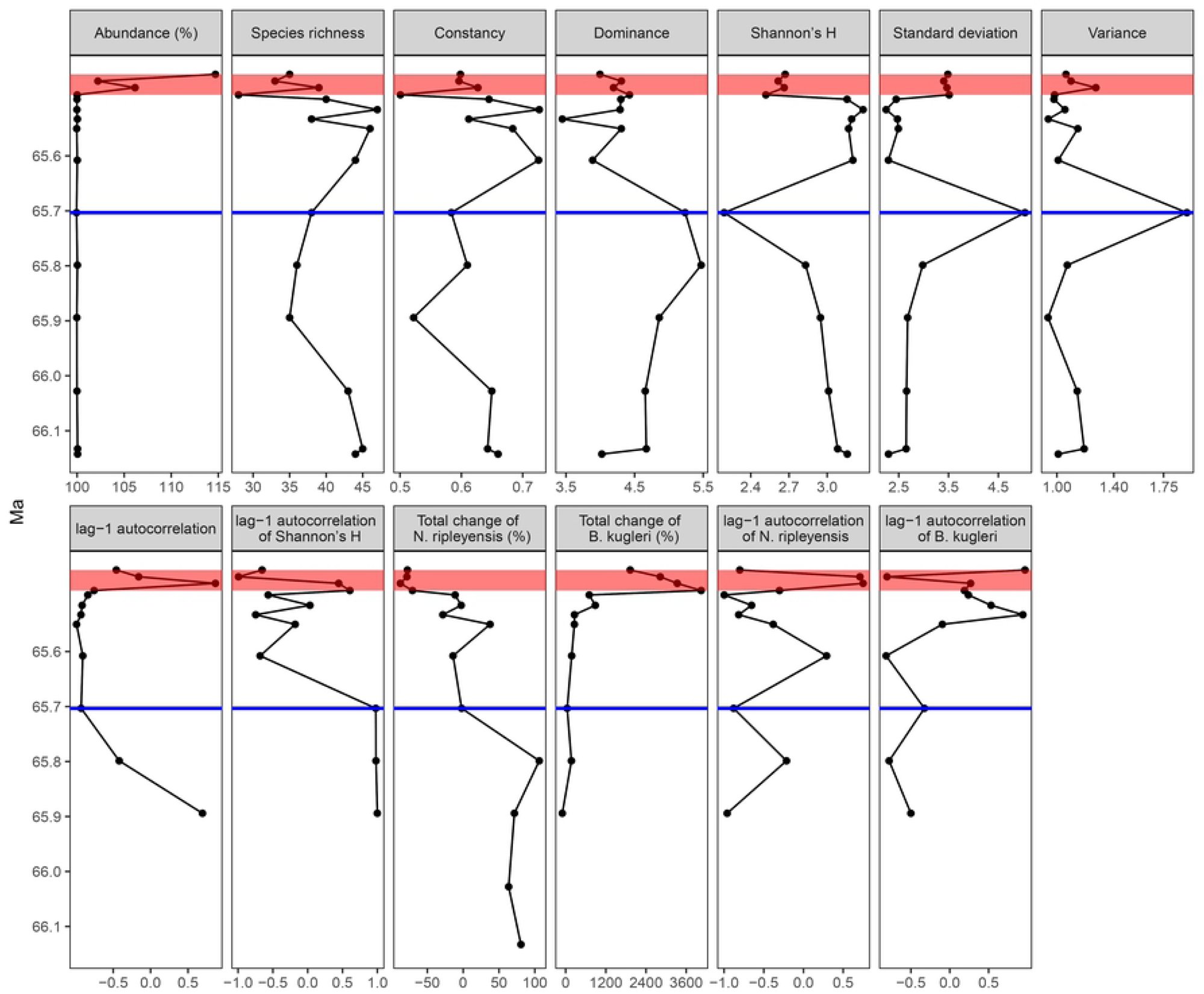
Collapse indicators of KPG data series in the pre-collapse and the collapse zones. Blue solid line = environmental event (bolide event), red box = collapse zone. Pre-collapse dominant species = *Nuttallinella ripleyensis*. Collapse dominant species = *Bulimina kugleri*. Source of data: Alegret and Thomas [21].

**Fig 12.**
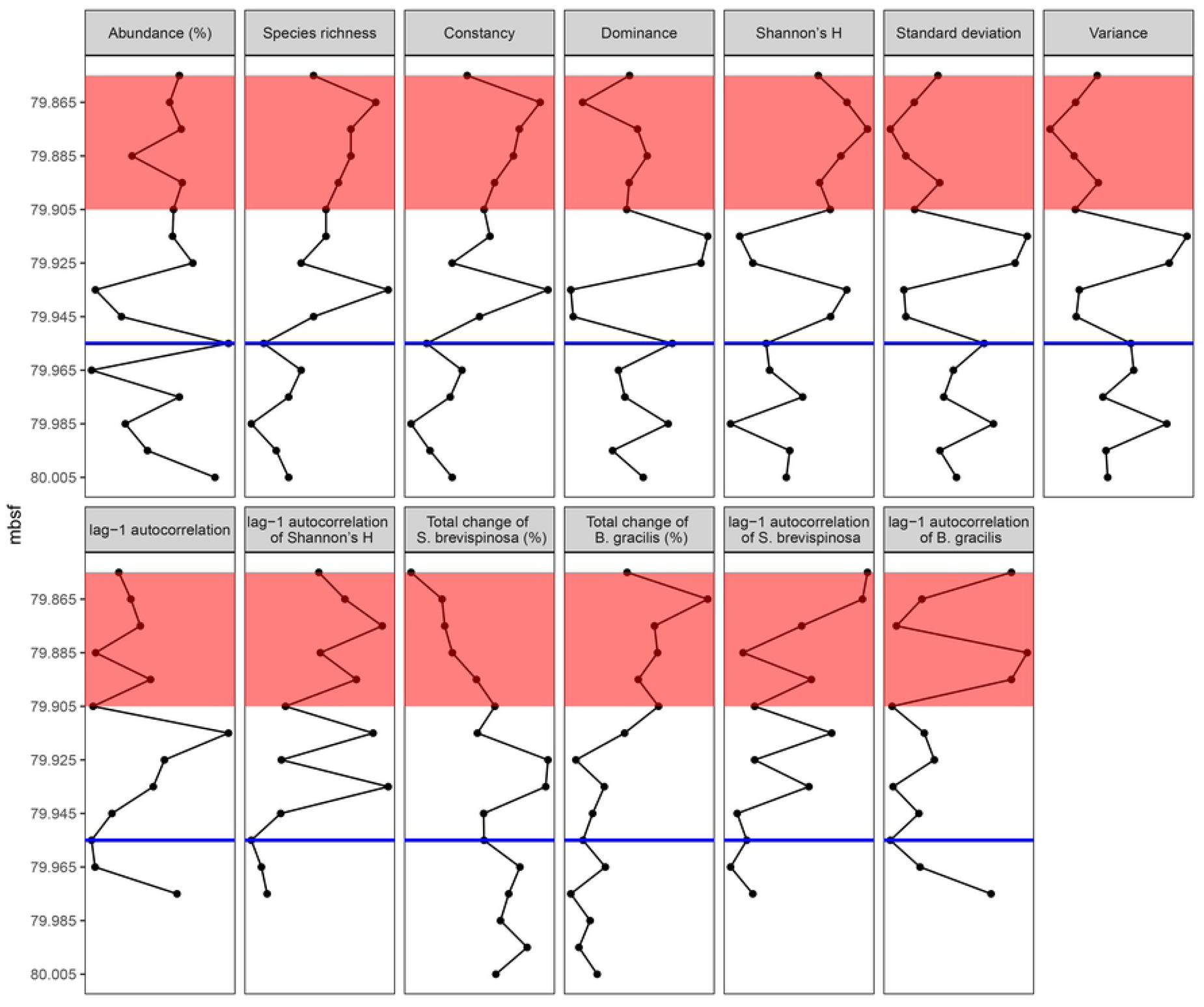
Collapse indicators of PE data series in the pre-collapse and collapse zones. Blue solid line = environmental event (assumed pulse event, temporary warming), red box = collapse zone. Pre-collapse dominant species= *Siphogenerinoides brevispinosa*. Collapse dominant species = *Bolivina gracilis*. Source of data: Takeda and Kaiho [23].

**Fig 13.**
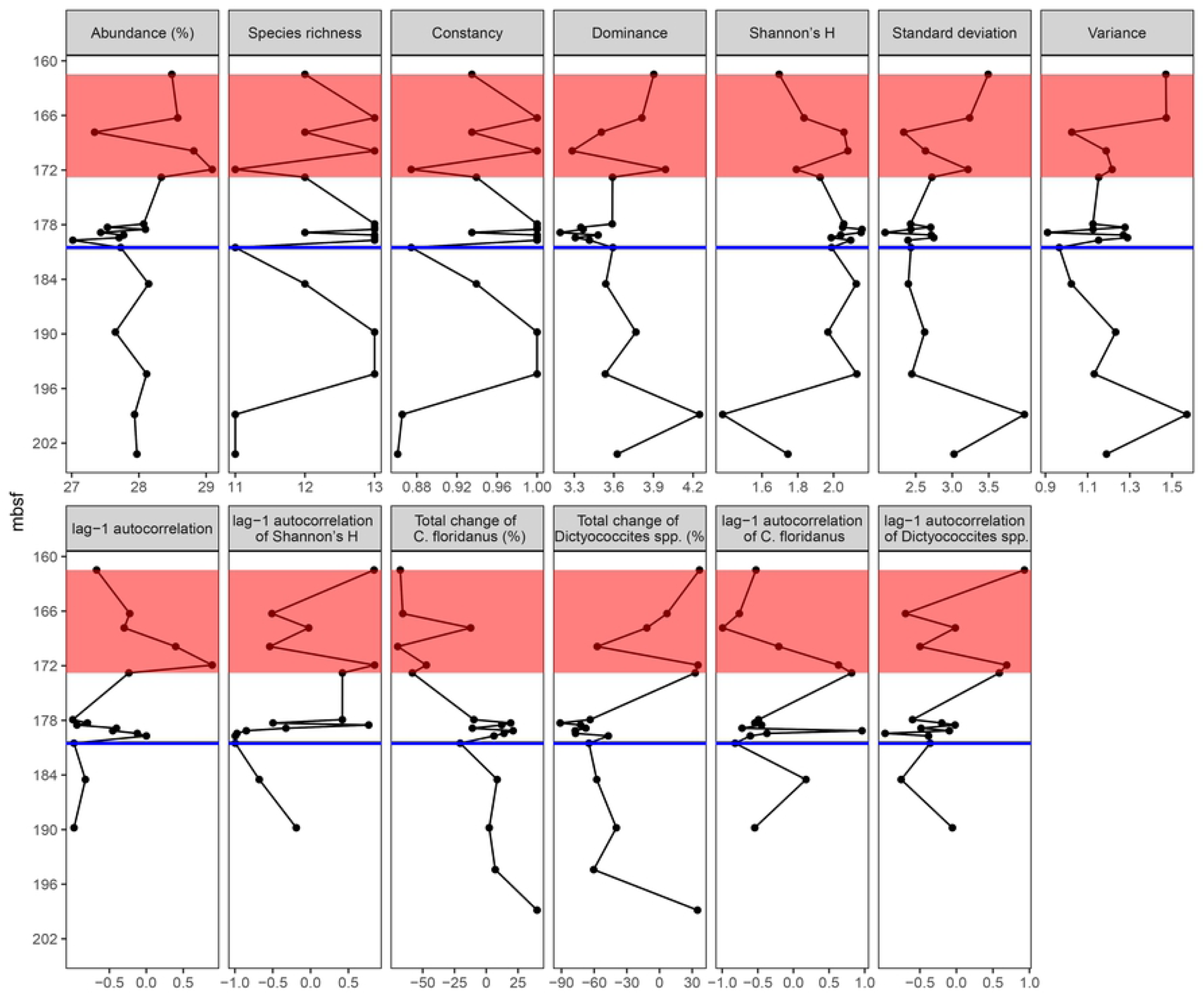
Collapse indicators of EM1 data series in the pre-collapse and collapse zones. Blue solid line = environmental event (temporary glaciation event), red box = collapse zone. Pre-collapse dominant species = *Cyclicargolithus floridanus*. Collapse dominant species = *Dictyococcites spp*. Source of data: Plancq et al. [24].

**Fig 14.**
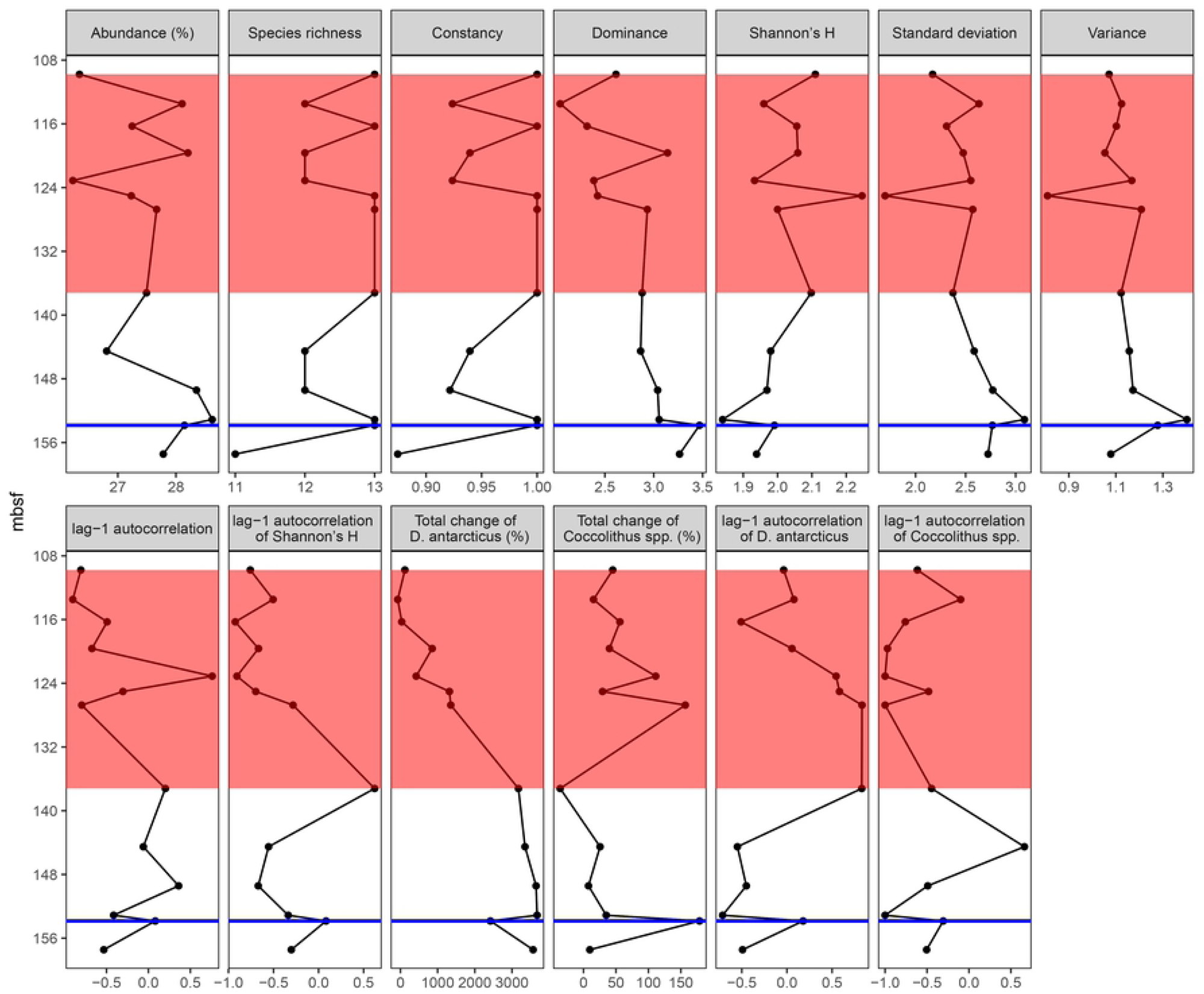
Collapse indicators of EM2 data series in the pre-collapse and the collapse zones. Blue solid line = environmental event (presumed temporary warming event), red box = collapse zone. Pre-collapse dominant species = *Dictyococcites antarcticus*. Collapse dominant species = *Coccolithus spp*. Source of data: Plancq et al. [24].

**Fig 15.**
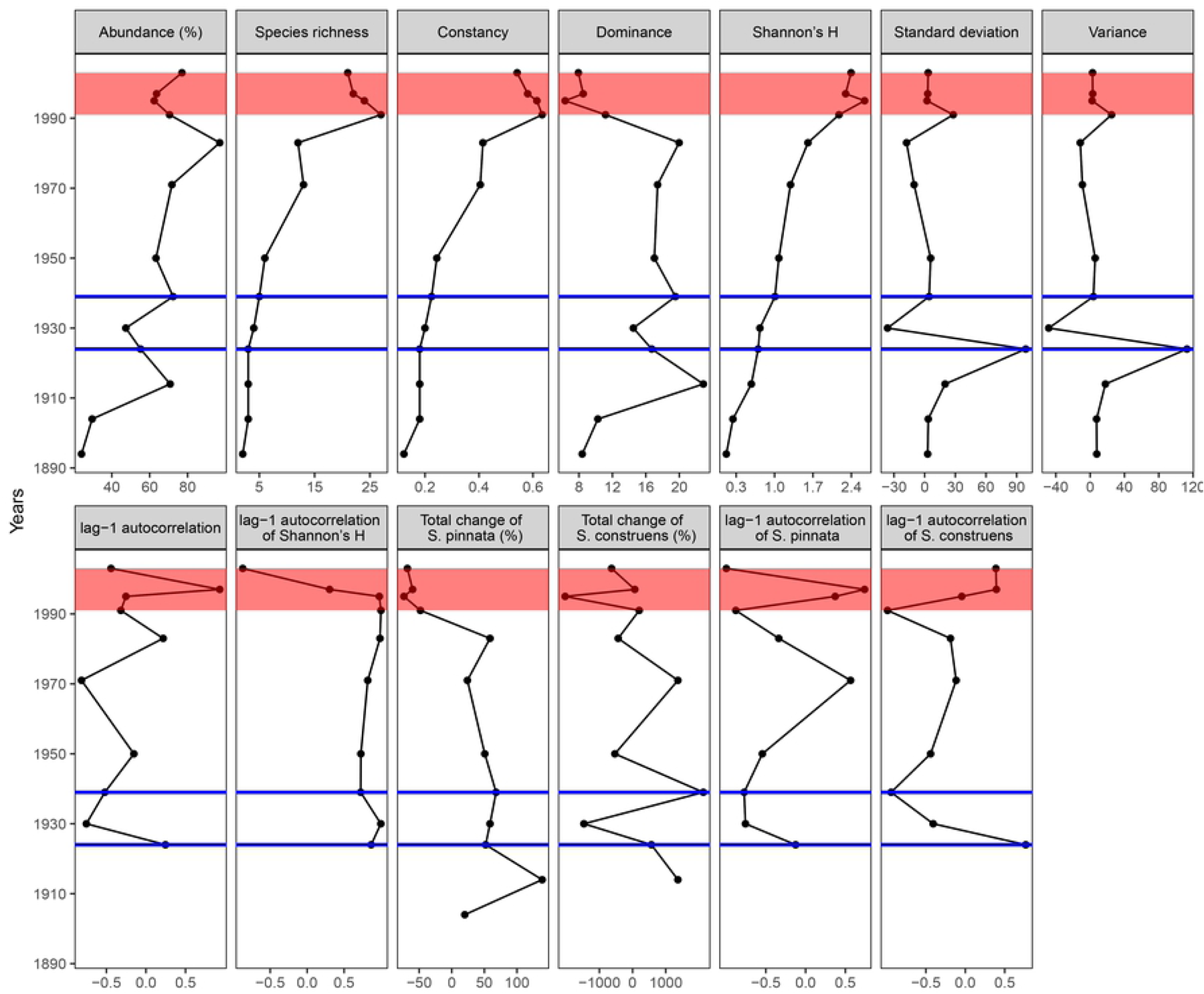
Collapse indicators of modern data series in the pre-collapse and the collapse zones. Blue solid line = environmental event (summer air temperature increase in 1924 and 1939), red box = collapse zone. Pre-collapse dominant species = *Staurosirella pinnata*. Collapse dominant species = *Staurosira construens*. Source of data: Köck et al. [25], Lehnherr et al. [20].

We tested the collapse indicators at different time scales, in different units, on different data types to detect universal signals. The time steps of the data series either in years or in mbsf (meters below seafloor). The time scales are in years, million years (Ma). In the evaluation of the indicators, we applied non-standard uniform units for the time steps. The probable collapse triggering environmental event is time step zero. Subsequent time steps are time steps 1, 2, 3, etc. in non-standard uniform units in each data series (KPG, PE, EM1, EM2, modern). Concerning the signals of the collapses, we make a difference between ‘small-scale signs’ and ‘large-scale signs’. Small-scale collapse signals are short, sharp signs (sharp drop or increase) at the environmental events or right after the events (or with a short lag). Small-scale signals are at the same time as large-scale signals, or they precede large-scale signals, sometimes with a gap. Small-scale signals are often earlier than large-scale signals. They indicate environmental events. Large-scale collapse signals are usually large-scale trend changes in the indicators. Large-scale signals are the signs of collapses.

We did not involve the modern data series into the evaluation of small-scale signals, because some indicators (abundance, dominance, standard deviation) give inconsistent signals at the environmental events in 1924 and 1939, respectively. Besides, we cannot explain the outlier at time step 1914. However, we must note that species richness, constancy, and variance increase suddenly after 1924. Shannon’s H also shows a sharper increase in 1939. We detected most small-scale signals after the probable environmental event in 1924. Large-scale signals appear after the event in 1939 indicated by our own-developed indicators (lag-1 autocorrelation of Shannon’s H, the total change of pre-collapse dominant species, lag-1 autocorrelation of pre-collapse dominant species). Hence, only the environmental event in 1939 is shown in the tables below as time step zero (Tables 1 and 2).

**Table 1.**
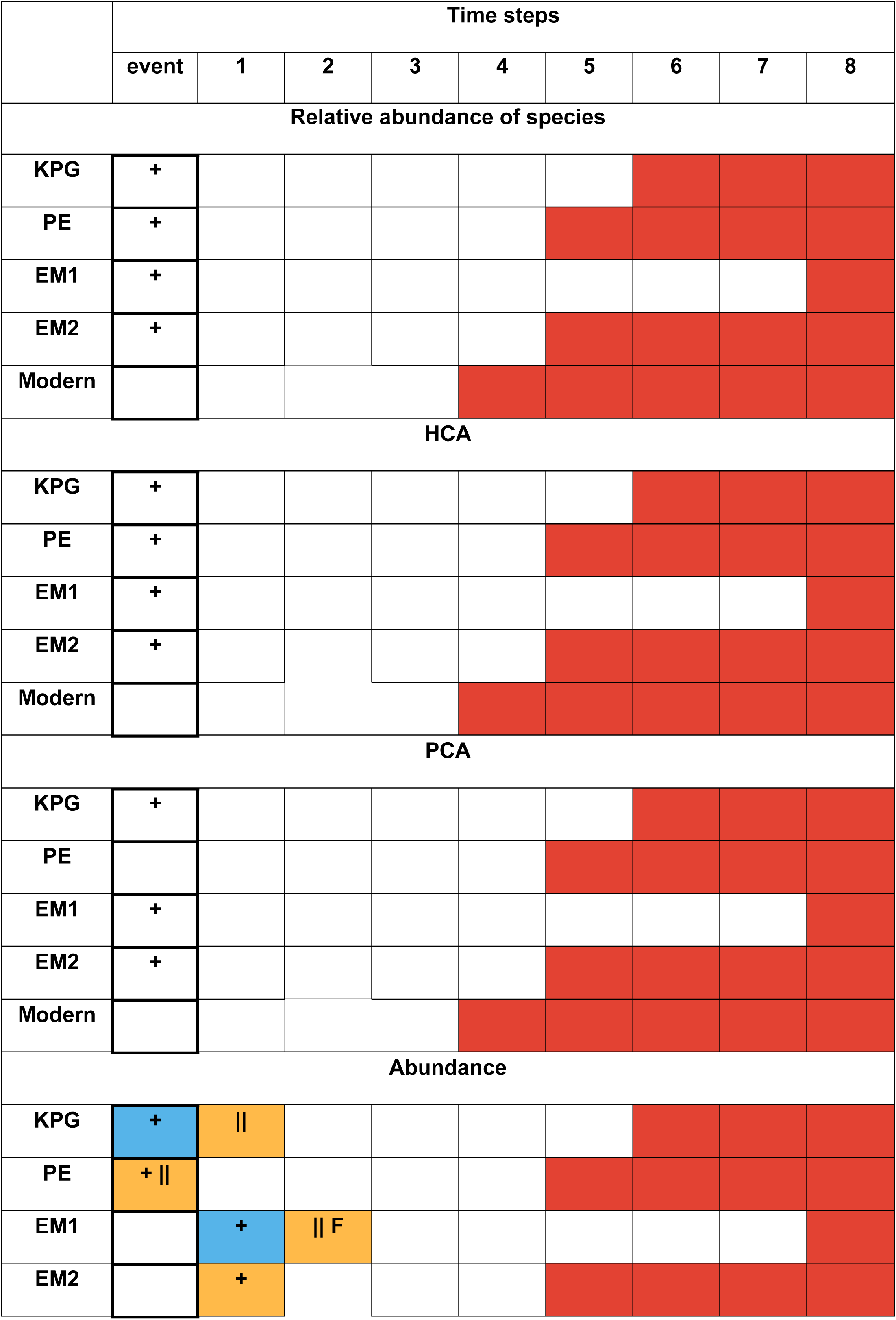
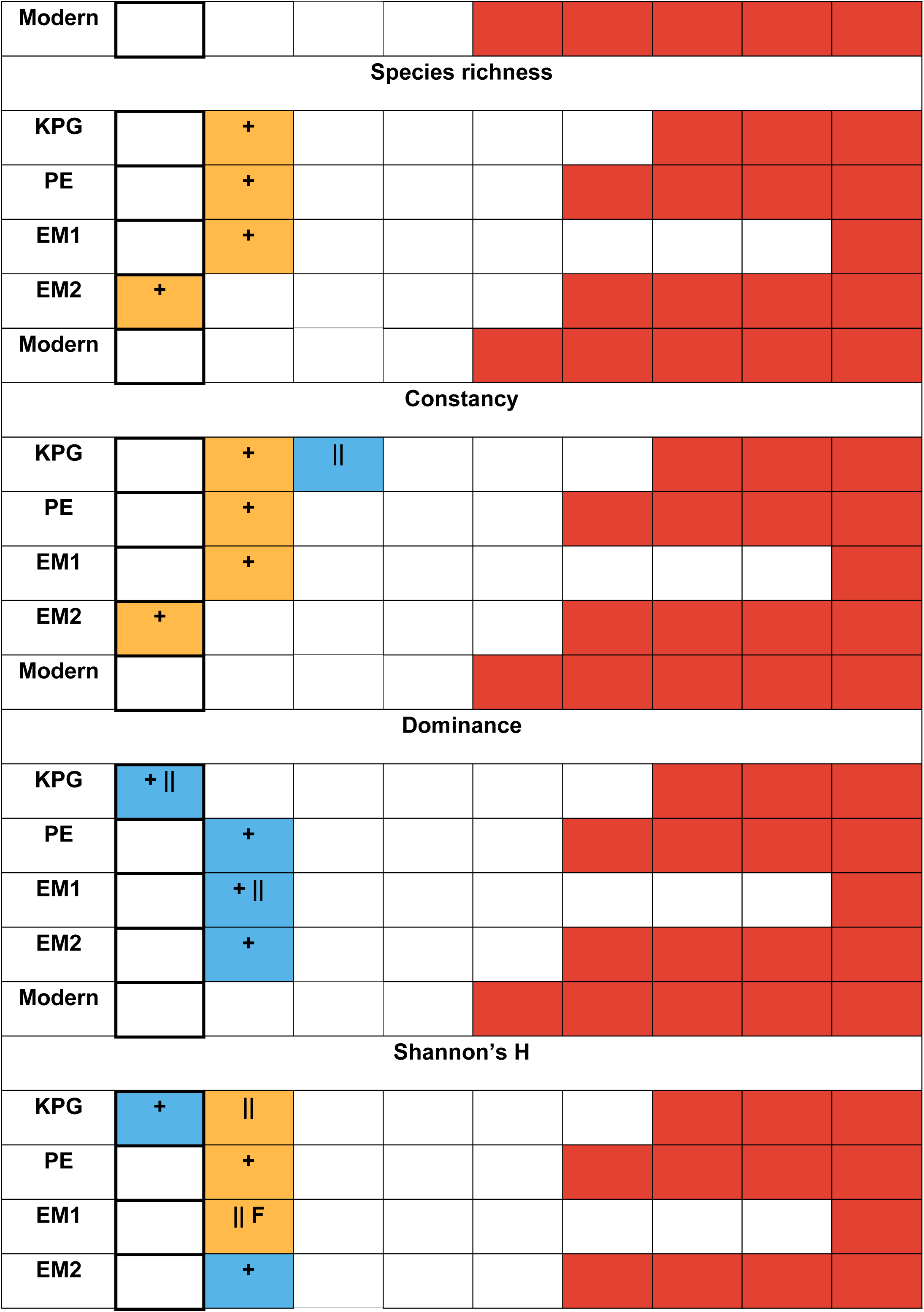
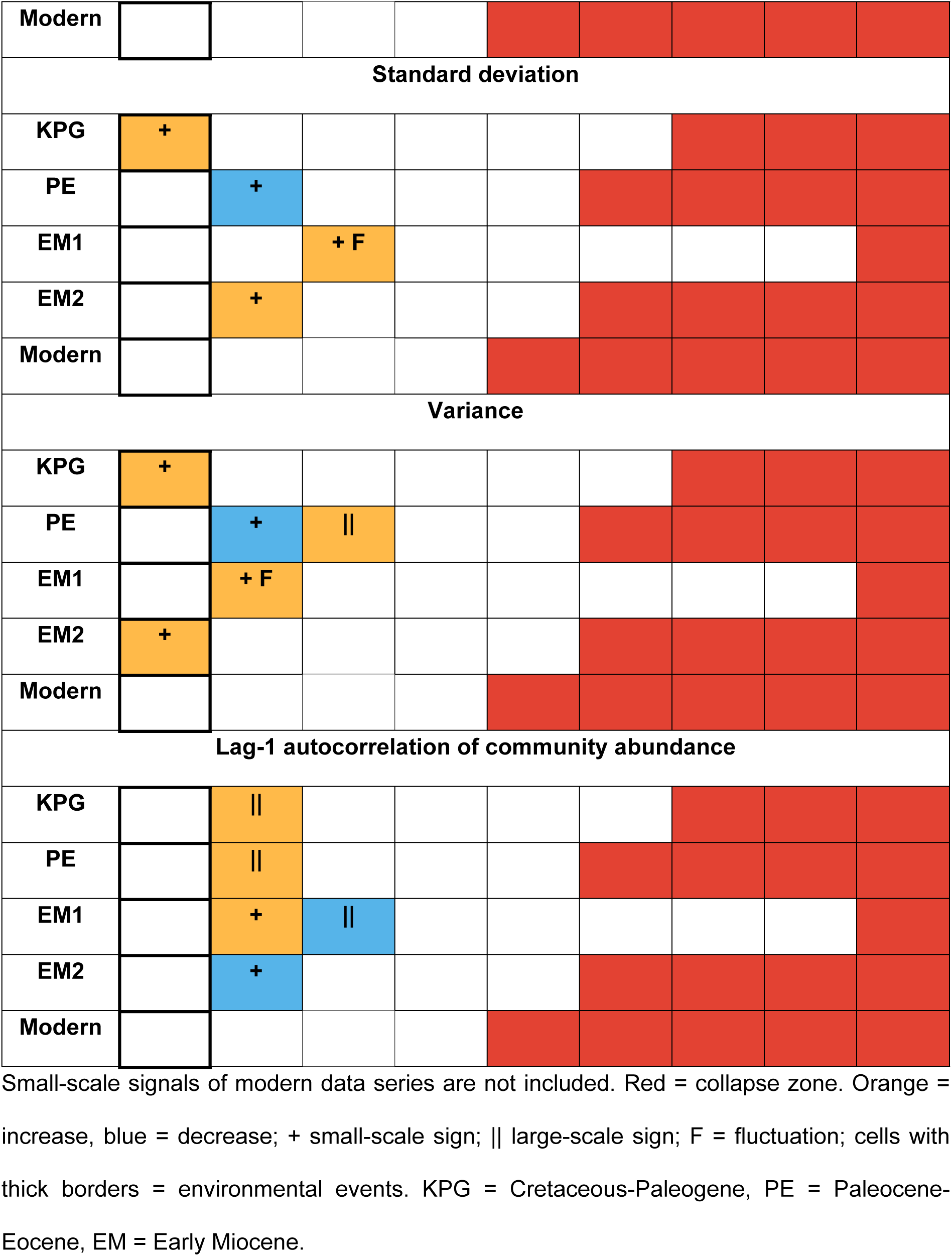
Small-scale and large-scale warning signals by indicators (relative abundance of species, HCA, PCA, abundance, species richness, constancy, dominance, Shannon’s H, standard deviation, variance, lag-1 autocorrelation) at community level.

**Table 2.**
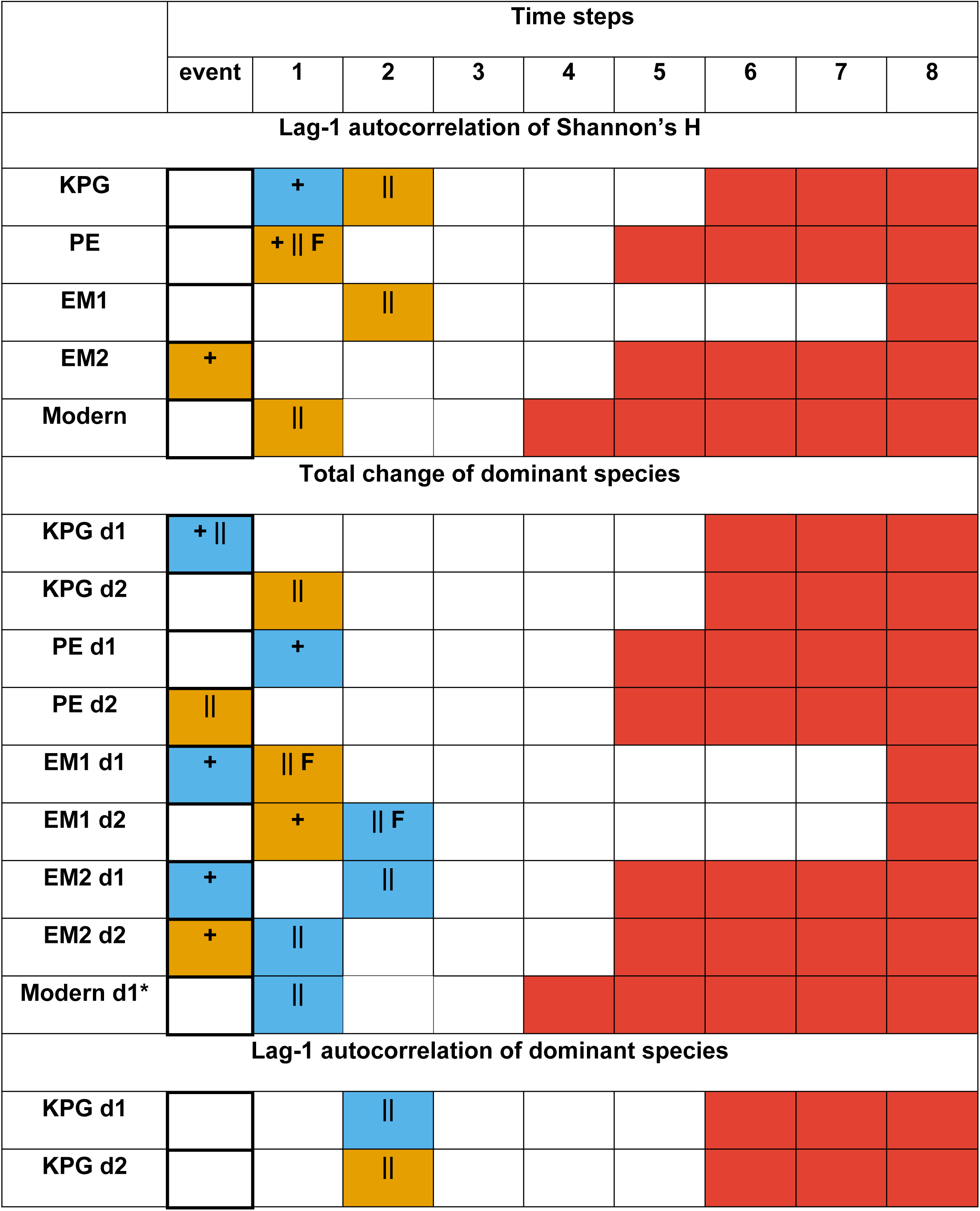
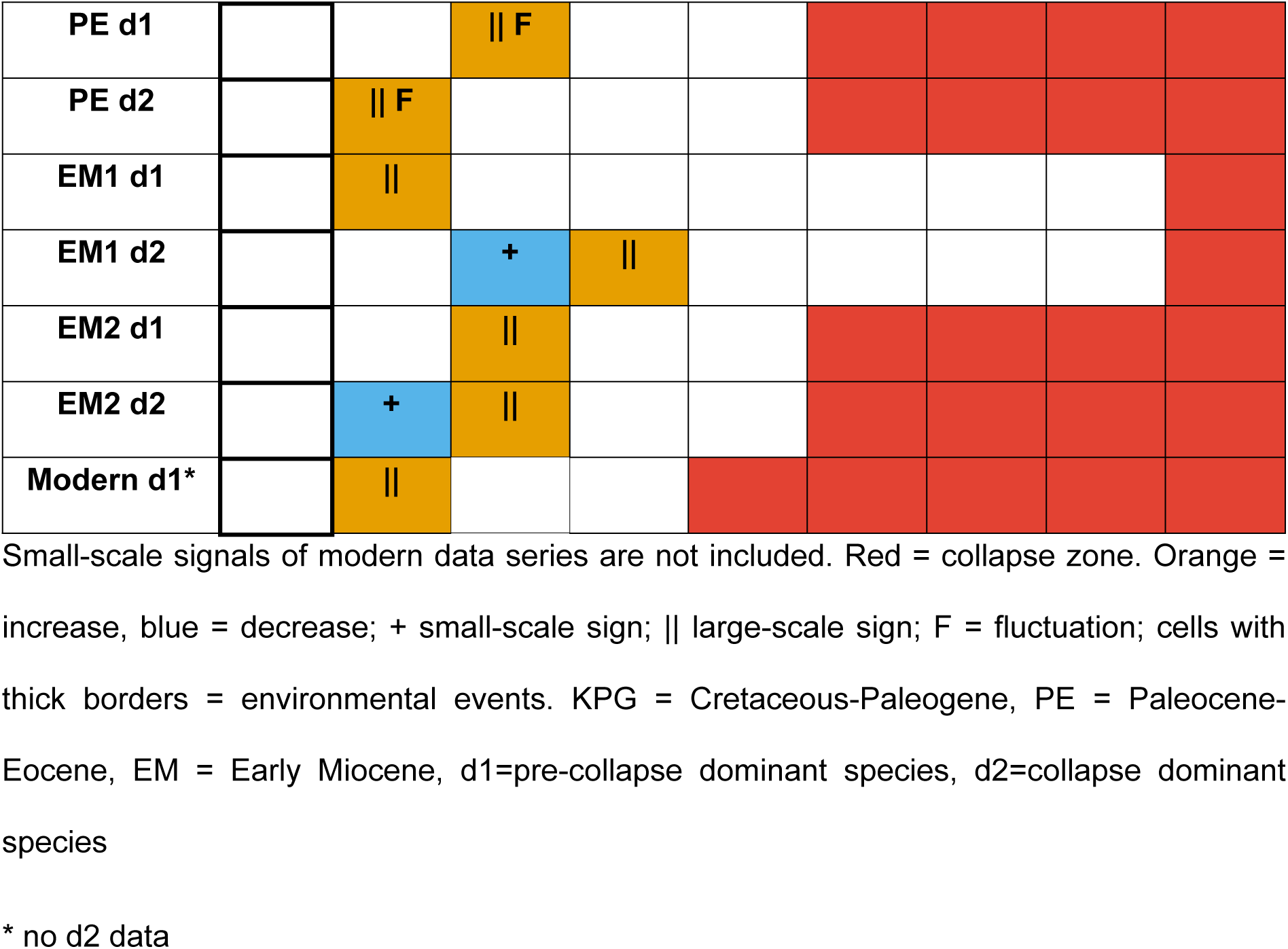
Small-scale and large-scale warning signals by own-developed indicators (lag-1 autocorrelation of Shannon’s H, the total change of pre-collapse and collapse dominant species, lag-1 autocorrelation of pre-collapse and collapse dominant species).

We evaluated the indicators by the number of signs they give close to the event and by their earliness (Tables 3–8). We also indicated if an indicator give consistent signals (increase or decrease). Small-scale signals are sharp decreases or increases at the environmental events. In the assessment of small-scale signals, we did not include the modern data series because of inconsistent signals at the environmental events in 1924 and 1939, respective.

**Table 3.** Ranking of indicators by the number of small-scale warning signs.

| Indicators | Number of small-scale signs | Ranking |
| --- | --- | --- |
| Abundance | 4 | 1 |
| Species richness | 4 | 1 |
| Constancy | 4 | 1 |
| Dominance | 4 | 1 |
| Standard deviation | 4 | 1 |
| Variance | 4 | 1 |
| Total change of pre-collapse dominant species | 4 | 1 |
| Combined indicator of the total change of pre-collapse and collapse dominant species | 4 | 1 |
| Relative abundance of species | 4 | 1 |
| **HCA** | **4** | **1** |
| **Shannon's H** | **3** | **2** |
| **Lag-1 autocorrelation of Shannon's H** | **3** | **2** |
| **PCA** | **3** | **2** |
| **Lag-1 autocorrelation of community abundance** | **2** | **3** |
| **Total change of collapse dominant species** | **2** | **3** |
| **Lag-1 autocorrelation of collapse dominant species** | **2** | **3** |
| **Combined indicator of lag-1 autocorrelation of pre-collapse and collapse dominant species** | **2** | **3** |
| **Lag-1 autocorrelation of pre-collapse dominant species** | **0** | **4** |

Large-scale signals are the signs of collapses and they are usually large-scale trend changes of the indicators. In the case of large-scale signs, we ignored indicators changing trends several times between the environmental event and the collapse zone boundary. We preferred indicators changing a trend close to the environmental event and then having an unchanged trend until / close to the collapse. To enhance the efficiency of indicators of the dominant species, we created a combined indicator of the total changes of the pre-collapse and collapse dominant species, as well as, that of the lag-1 autocorrelation of the pre-collapse and the collapse dominant species. It means that we considered either the sign of the pre-collapse dominant species or the collapse dominant species during the evaluation, depending on their earliness.

We identified a collapse-warning zone based on the detected signals. The collapse-warning zone starts at the environmental event and it includes the small-scale and large-scale signals. It is shorter than the whole pre-collapse period. In this study, the warning signal zones consist of three or four time steps (the environmental event plus two or three time steps).

First, we evaluated the indicators by the number of signs the indicators give close to the event (Tables 3 and 4). The indicators with the highest number of signals get the best ranks. The small-scale signals of the modern data series are not included.

**Table 4.** Ranking of indicators by the number of large-scale warning signs.

| **Indicators** | **Number of large-scale signs** | **Ranking** |
| --- | --- | --- |
| **Combined indicator of the total change of pre-collapse and collapse dominant species** | **5** | **1** |
| **Lag-1 autocorrelation of pre-collapse dominant species** | **5** | **1** |
| **Combined indicator of lag-1 autocorrelation of pre-collapse and collapse dominant species** | **5** | **1** |
| **Lag-1 autocorrelation of Shannon's H** | **4** | **2** |
| **Total change of pre-collapse dominant species** | **4** | **2** |
| **Total change of collapse dominant species** | **4** | **2** |
| **Lag-1 autocorrelation of collapse dominant species** | **4** | **2** |
| **Abundance** | **3** | **3** |
| **Lag-1 autocorrelation of community abundance** | 3 | 3 |
| **Dominance** | 2 | 4 |
| **Shannon's H** | 2 | 4 |
| **Variance** | 2 | 4 |
| **Constancy** | 1 | 5 |
| **Standard deviation** | 1 | 5 |
| **Species richness** | 0 | 6 |

Second, we evaluated the signals by their earliness (Tables 5 and 6). We detected the signals of the collapse warning zones between time steps 0–3 (time step zero = environmental event). Based on the earliness, the time steps get weights as follows: event = 4; step1 = 3; step2 = 2; step3 = 1. The indicators with the highest weight of signals get the best ranks. The small-scale signals of the modern data series are not included.

**Table 5.** Ranking of indicators by the earliness of small-scale warning signs.

| **Indicators** | **Weight of**<br>**small-scale**<br>**signs** | **Ranking** |
| --- | --- | --- |
| **Relative abundance of species** | 12 | 1 |
| **HCA** | 12 | 1 |
| **Total change of pre-collapse dominant species** | 11 | 2 |
| **Combined indicator of the total change of pre-collapse**<br>**and collapse dominant species** | 11 | 2 |
| **Abundance** | 10 | 3 |
| **Variance** | 10 | 3 |
| **Species richness** | 9 | 4 |
| **Constancy** | 9 | 4 |
| **Dominance** | 9 | 4 |
| **PCA** | 9 | 4 |
| **Standard deviation** | 8 | 5 |
| **Shannon's H** | 7 | 6 |
| **Lag-1 autocorrelation of Shannon's H** | 7 | 6 |
| **Total change of collapse dominant species** | 5 | 7 |
| **Lag-1 autocorrelation of community abundance** | 4 | 8 |
| **Lag-1 autocorrelation of collapse dominant species** | 3 | 9 |
| **Combined indicator of lag-1 autocorrelation of pre-collapse and collapse dominant species** | 3 | 9 |
| **Lag-1 autocorrelation of pre-collapse dominant species** | 0 | 10 |

**Table 6.** Ranking of indicators by the earliness of large-scale signs.

| **Indicators** | **Weight of large-scale signs** | **Ranking** |
| --- | --- | --- |
| **Combined indicator of the total change of pre-collapse and collapse dominant species** | 15 | 1 |
| **Total change of collapse dominant species** | 12 | 2 |
| **Combined indicator of lag-1 autocorrelation of pre-collapse and collapse dominant species** | 11 | 3 |
| **Lag-1 autocorrelation of pre-collapse dominant species** | 10 | 4 |
| **Abundance** | 9 | 5 |
| **Total change of pre-collapse dominant species** | 9 | 5 |
| **Lag-1 autocorrelation of community abundance** | 8 | 6 |
| **Lag-1 autocorrelation of Shannon's H** | 8 | 6 |
| **Lag-1 autocorrelation of collapse dominant species** | 8 | 6 |
| **Dominance** | 7 | 7 |
| **Shannon's H** | 6 | 8 |
| **Variance** | 5 | 9 |
| **Constancy** | 2 | 10 |
| **Standard deviation** | 2 | 10 |
| **Species richness** | 0 | 11 |

To get the overall ranking of indicators, we summed up the ranks of indicators based on the number and the earliness of signs (table 7–8). The best-performing indicators have the smallest sum. We also indicated the consistency of signs (consistent decrease or increase of indicators in all data series).

**Table 7.** Ranking of indicators based on the number and the earliness of small-scale signals.

| **Indicators** | **Ranking** | **Consistency** |
| --- | --- | --- |
| **Relative abundance of species** | 1 | No |
| **HCA** | 1 | No |
| **Total change of pre-collapse dominant species** | 2 | Yes<br>(decrease) |
| **Combined indicator of the total change of pre-collapse**<br>**and collapse dominant species** | 2 | No |
| **Abundance** | 3 | No |
| **Variance** | 3 | No |
| **Species richness** | 4 | Yes<br>(increase) |
| **Constancy** | 4 | Yes<br>(increase) |
| **Dominance** | 4 | Yes<br>(decrease) |
| **Standard deviation** | 5 | No |
| **PCA** | 5 | No |
| **Shannon's H** | 6 | No |
| **Lag-1 autocorrelation of Shannon's H** | 6 | No |
| **Total change of collapse dominant species** | 7 | No |
| **Lag-1 autocorrelation of community abundance** | 8 | No |
| **Lag-1 autocorrelation of collapse dominant species** | 9 | No |
| **Combined indicator of lag-1 autocorrelation of pre-collapse and collapse dominant species** | 9 | No |
| **Lag-1 autocorrelation of pre-collapse dominant species** | 10 | No |

HCA along with the relative abundance of species are the best small-scale-signal indicators in this study (Tables 1, 3, 5 and 7). They indicate the environmental events the best. HCA shows unusual environmental events as an outlier precisely (Figs 3–5, Table 1). The relative abundance of the pre-collapse dominant species usually drops and the relative abundance of environmental indicators shoots up at unusual environmental events (Figs 3–8, Table 1). The total change of the pre-collapse dominant species and the combined indicator of the total change of pre-collapse and collapse dominant species are also good indicators (Figs 11–15, Tables 2 and 3, 5, 7). The total change of the pre-collapse dominant species gives a consistent sign, as it decreases suddenly at the environmental events. The abundance, variance, species richness, constancy, dominance also give small-scale signs in all data series, however, they are not the earliest indicators (Figs 11–15, Tables 1, 3, 5 and 7). Nevertheless, species richness, constancy, and dominance are good environmental event indicators, because their signals are consistent: species richness and constancy increase sharply, while dominance has a sudden drop. Indicators combined with lag-1 autocorrelation underperform as small-scale signal indicators in our analysis (Figs 11–15, Table 7)

The best-performing large-scale-signal indicators are the indicators of the dominant species based on the number and the earliness of signals (Table 8). The combined indicator of the total change of pre-collapse and collapse dominant species is the most effective collapse indicator, as it considers the signals of both pre-collapse and collapse dominant species. Community-level indicators lag behind. The abundance and the lag-1 autocorrelation of Shannon’s H are mediocre indicators, however, they give consistent signs: they start an increasing trend as the warning sign of the approaching collapse (Figs 11–15, Tables 1 and 2). Dominance also has a consistent collapse signal. It starts a decreasing trend at/after the environmental events (Figs 11–15, Table 1). In this study, the weakest collapse indicators are constancy, standard deviation, and species richness (Table 8). The rankings of indicators can change depending, for instance, on the number of data series involved in the analysis, however, the clusters of strong and weak indicators probably remain similar. The total change of the dominant species and the combined version of this indicator outperform community-level critical slowing down indicators (lag-1 autocorrelation of community abundance, variance, standard deviation) and give a warning signal much more reliably than any other indicators in this study.

**Table 8.**
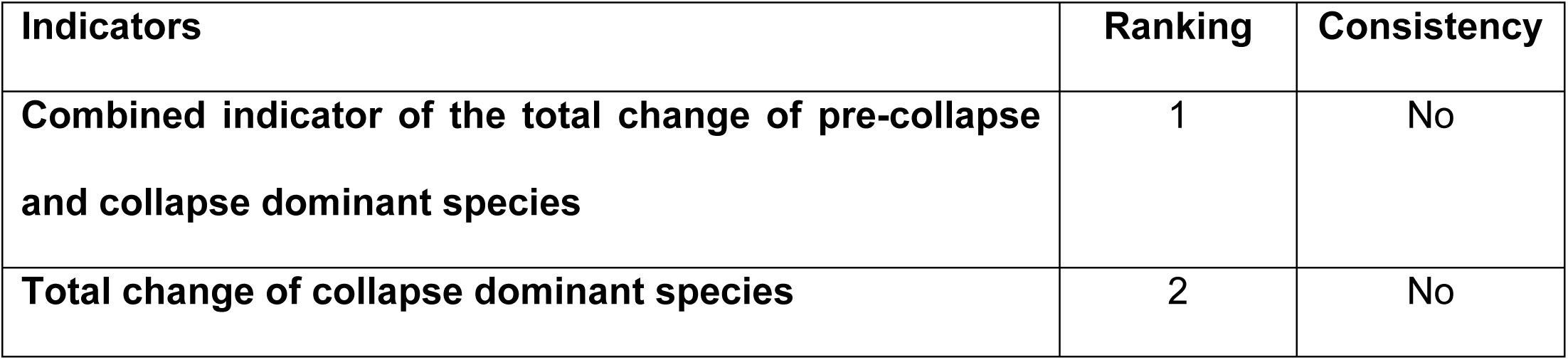
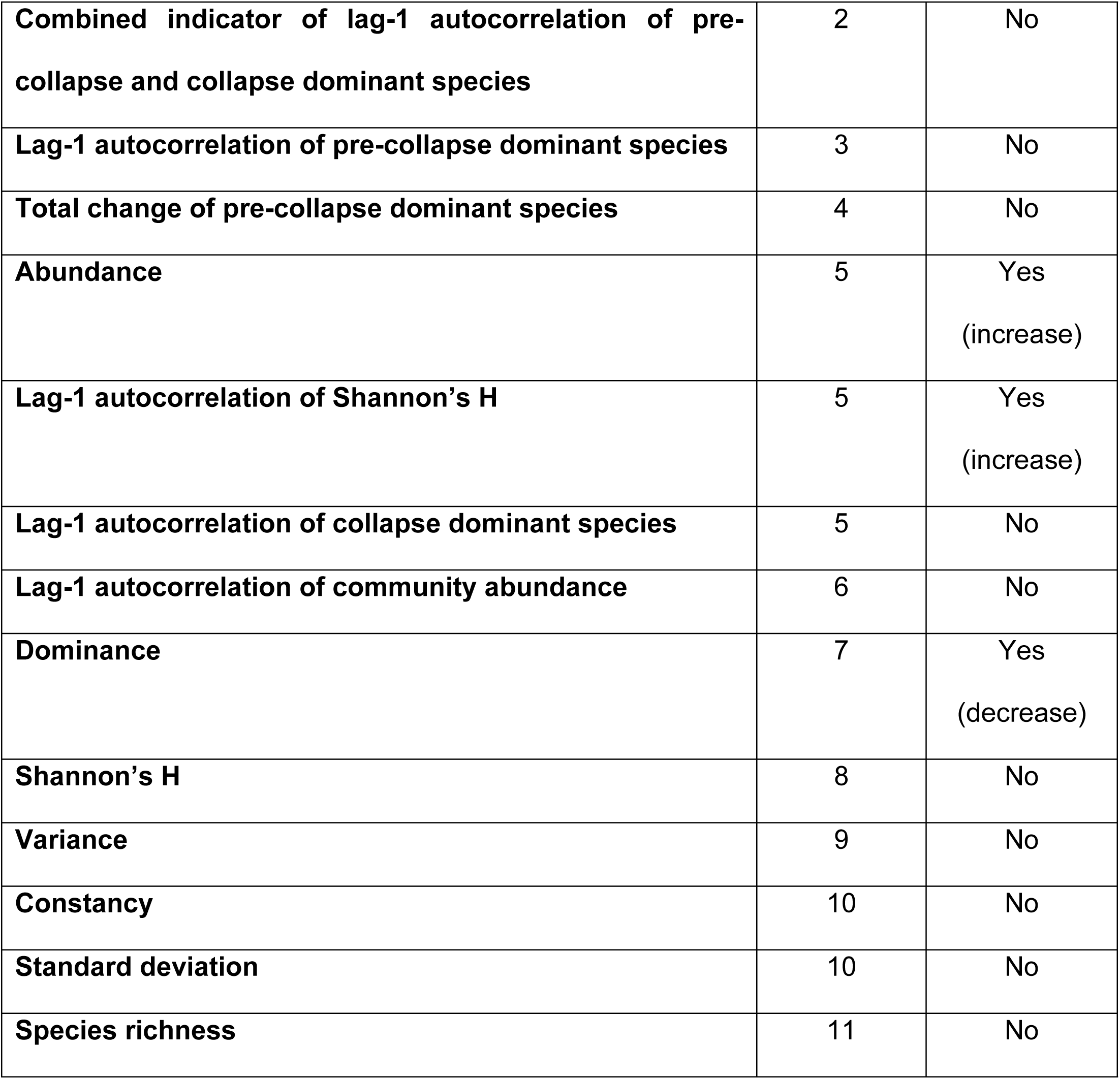
Ranking of indicators based on the number and the earliness of large-scale signals.

| **Indicators** | **Ranking** | **Consistency** |
| --- | --- | --- |
| **Combined indicator of the total change of pre-collapse and collapse dominant species** | 1 | No |
| **Total change of collapse dominant species** | 2 | No |
| **Combined indicator of lag-1 autocorrelation of pre-collapse and collapse dominant species** | 2 | No |
| **Lag-1 autocorrelation of pre-collapse dominant species** | 3 | No |
| **Total change of pre-collapse dominant species** | 4 | No |
| **Abundance** | 5 | Yes<br>(increase) |
| **Lag-1 autocorrelation of Shannon's H** | 5 | Yes<br>(increase) |
| **Lag-1 autocorrelation of collapse dominant species** | 5 | No |
| **Lag-1 autocorrelation of community abundance** | 6 | No |
| **Dominance** | 7 | Yes<br>(decrease) |
| **Shannon's H** | 8 | No |
| **Variance** | 9 | No |
| **Constancy** | 10 | No |
| **Standard deviation** | 10 | No |
| **Species richness** | 11 | No |

To sum it up, small-scale signs indicate collapse-triggering environmental events. HCA usually has distinctive outliers at the environmental events. PCA can also have an outlier at the environmental events. Other indicators have a sharp drop or a shoot up as a small-scale sign.

The relative abundance of environmental indicators increases sharply, the relative abundance of pre-collapse dominant species drops sharply at the environmental events. The HCA and the relative abundance of species together are reliable indicators of the environmental events. It seems to be a general rule, at least, based on the results of this study that the species richness and the constancy increase suddenly as a small-scale sign at the environmental events. The total change of the pre-collapse dominant species and the dominance drop; the total change of the collapse species increases suddenly (when it gives a signal) as a small-scale sign at the environmental events in the pre-collapse period. The indicators of the dominant species are the best large-scale indicators. They outperform the indicators of critical slowing down (standard deviation, variance, lag-1 autocorrelation of community abundance) in this study. Our development, the combined indicator of the total change of pre-collapse and collapse dominant species is the most reliable collapse indicator. Our other development, the lag-1 autocorrelation of Shannon’s H is a mediocre indicator based on the evaluation of this study, however, it usually gives a very distinctive sign, hence it is worth further testing. Considering variance and standard deviation, variance is a better indicator, however, neither of them is a reliable and consistent indicator.

## Conclusions

In this study, one of our objectives was to test and develop collapse indicators. We involved traditional community indicators (HCA, PCA, relative abundance of species, abundance, species richness, constancy, dominance, Shannon’s H) and CSD indicators (standard deviation, variance, lag-1 autocorrelation) into the analysis. We also developed/further developed indicators. These indicators are the lag-1 autocorrelation of Shannon’s H, the total change of dominant species and the lag-1 autocorrelation of dominant species.

We distinguished between two groups of warning signals: small-scale and large-scale warning signals. Small-scale signs indicate collapse triggering environmental events. Outliers of HCA and PCA, sharp peaks or drops of other indicators are small-scale signs. Large-scale collapse signals are usually large-scale trend changes of the indicators between the environmental event and the collapse boundary. They forecast the collapses.

We evaluated the indicators by the number and the earliness of the warning signs they give. The best small-scale-signal indicators are the HCA and the relative abundance of species. Our development, the total change of the pre-collapse dominant species and the combined indicator of the total change of pre-collapse and collapse dominant species are also good small-scale-signal indicators. The total change of the pre-collapse dominant species, species richness, constancy, and dominance give consistent small-scale signs. The total change of the pre-collapse dominant species and dominance usually decrease, while species richness and constancy decrease at/after unusual environmental events. Indicators combined with lag-1 autocorrelation underperform as environmental event indicators.

The best large-scale-sign indicators are the indicators of the dominant species. Our development, the combined indicator of the total change of pre-collapse and collapse dominant species is the most effective collapse indicator. Community-level collapse indicators lag behind. An advantage of abundance, lag-1 autocorrelation of Shannon’s H and dominance that they give a consistent large-scale sign. Abundance and lag-1 autocorrelation of Shannon’s H start an increasing trend, whereas dominance starts a decreasing trend as a collapse-warning signal.

Considering both small-scale and large-scale warning signals, our new indicator, the combined indicator of the total change of pre-collapse and collapse dominant species is the best collapse indicator among the studied collapse indicators. This finding supports our hypothesis that dominant species (in this study, the most abundant species) are sensitive to unusual environmental events and their decline is a main contributor to community collapses. The efficiency of our combined indicator might also refer to the fact that the rise of the collapse community starts close to the collapse-triggering environmental event. Our other development, the lag-1 autocorrelation of Shannon’s H, gives one of the most distinctive signs, however, based on the number and the earliness of the signs, it is a mediocre large-scale collapse indicator.

In this study, we showed that unusual environmental events may be the number one environmental triggers of community collapses. Dominant species (the most abundant species) are sensitive to environmental events and they start to decline immediately as a response to the environmental event. The decline of the pre-collapse dominant species may be an important biotic (secondary) cause of collapses. The combined indicator of the total change of pre-collapse and collapse dominant species proved to be an effective small-scale and large-scale collapse indicator. Small-scale signals should be involved in studies because they can be earlier than large-scale trend changes of collapse indicators.

## Abbreviations of periods, epochs and events

KPG: Cretaceous-Paleogene
PE: Paleocene-Eocene
PETM: Paleocene-Eocene Thermal Maximum
EM1: Early Miocene 1
EM2: Early Miocene 2

## Acknowledgments

We thank Dr. John P. Smol (Queen’s University, Kingston, Canada), Dr. Neal Michelutti (Queen’s University, Kingston, Canada) and Dr. Igor Lehnherr (University of Toronto- Mississauga, Mississauga, Canada) for providing diatom data from Lake Hazen.

## Supporting information

**S1 Appendix. Deficiencies and uncertainties of data series.**

